# Unraveling the genetic architecture of grain size in einkorn wheat through linkage and homology mapping, and transcriptomic profiling

**DOI:** 10.1101/377820

**Authors:** Kang Yu, Dongcheng Liu, Yong Chen, Dongzhi Wang, Wenlong Yang, Wei Yang, Lixin Yin, Chi Zhang, Shancen Zhao, Jiazhu Sun, Chunming Liu, Aimin Zhang

## Abstract

Genome-wide linkage and homology mapping revealed 17 genomic regions through a high-density einkorn wheat genetic map constructed using RAD-seq, and transcription levels of 20 candidate genes were explored using RNA-seq.

**Abstract:** Understanding the genetic architecture of grain size is a prerequisite to manipulate the grain development and improve the yield potential in crops. In this study, we conducted a whole genome-wide QTL mapping of grain size related traits in einkorn wheat by constructing a high-density genetic map, and explored the candidate genes underlying QTL through homologous analysis and RNA sequencing. The high-density genetic map spanned 1873 cM and contained 9937 SNP markers assigned to 1551 bins in seven chromosomes. Strong collinearity and high genome coverage of this map were revealed with the physical maps of wheat and barley. Six grain size related traits were surveyed in five agro-climatic environments with 80% or more broad-sense heritability. In total, 42 QTL were identified and assigned to 17 genomic regions on six chromosomes and accounted for 52.3-66.7% of the phenotypic variations. Thirty homologous genes involved in grain development were located in 12 regions. RNA sequencing provided 4959 genes differentially expressed between the two parents. Twenty differentially expressed genes involved in grain size development and starch biosynthesis were mapped to nine regions that contained 26 QTL, indicating that the starch biosynthesis pathway played a vital role on grain development in einkorn wheat. This study provides new insights into the genetic architecture of grain size in einkorn wheat, the underlying genes enables the understanding of grain development and wheat genetic improvement, and the map facilitates the mapping of quantitative traits, map-based cloning, genome assembling and comparative genomics in wheat taxa.

## Introduction

Grain weight is one of the most important traits in wheat (*Triticum aestivum* L.), which were mainly and tightly underpinned by grain morphology including two main components, grain length and width. In domestication process and breeding history, grain size was a major selection and breeding target, and has been widely selected and manipulated to increase the related grain yield in wheat (Gegas *et al.*, 2010). In China, an increase in wheat yield potential from ∼1 T ha^-1^ in 1965 to ∼5.4 T ha^-1^ today is mainly due to the great genetic increase in thousand-grain weight (NBS, 2015). Meanwhile, the grain morphology directly influences the milling performance and seedling vigor, which in turn determines the end products (Campbell *et al.*, 1999; Gegas *et al.*, 2010). Moreover, wheat preserves huge variations of grain size and weight among domesticated and wild species in diploid, tetraploid and hexaploid levels (Gegas *et al.*, 2010; Jing *et al.*, 2007). Thus, understanding the genetic structures of grain size would provide prerequisite information for wheat improvements.

High-density genetic maps played a fundamental role in dissecting genetic components of agronomic traits and assembling genomes. As a basic tool in genetic and genomic researches, high-density genetic maps have been widely developed in crops, including cereal crops, e.g., wheat (Iehisa *et al.*, 2014; Kumar *et al.*, 2016), rice (Xie *et al.*, 2010), maize (Chen *et al.*, 2014), and barley (Chutimanitsakun *et al.*, 2011), and economic crops, e.g., eggplant (Barchi *et al.*, 2012), grape (Wang *et al.*, 2012a), and sesame (Zhang *et al.*, 2013). A high-density consensus map of tetraploid wheat was developed by integrating datasets of 13 bi-parental populations, which harbored 30144 markers and covered 2631 cM of A and B sub-genomes (Maccaferri *et al.*, 2015). In hexaploid wheat, a high-density genetic map was constructed recently including 119566 single nucleotide polymorphism (SNP) markers, greatly facilitating the fine-mapping of a major QTL for grain number (Cui *et al.*, 2017). In barley, a high-density amplified fragment-length polymorphism map of 3H involving 84 markers covered 6.7 cM and was applied to narrow the genomic region for the important domestication loci, *Brittle rachis* (*Btr1* and *Btr2*), which have been further molecular cloned and characterized (Komatsuda *et al.*, 2004; Pourkheirandish *et al.*, 2015). Recently, with the demand of genome sequencing, the high-density genetic maps have been widely exploited at genome assembling, especially for the construction of chromosomal pseudomolecules by anchoring and ordering scaffolds, i.e. wheat (Chapman *et al.*, 2015; Jia *et al.*, 2013), cotton (Zhang *et al.*, 2015), and peanut (Bertioli *et al.*, 2016).

For the species with less available genomic information, it is urgently necessary to develop a high-density genetic map with a large number of genetic markers distributed over the whole genome. The advances in high-throughput sequencing technologies provide an excellent platform for genome-wide discovery of sequence variations and development of polymorphic genetic markers. Of these, genotyping-by-sequencing (GBS) methods utilize restriction enzyme digestion to reduce the complexity of a genome and sequence large amount of the resulted fragments with next-generation sequencing platforms, e.g., HiSeq 2000. This process would provide huge SNPs for the development of high-density genetic linkage map. Moreover, the methods, such as restriction site-associated DNA sequencing (RAD-seq) (Baird *et al.*, 2008), allow to label fragments with barcode sequences and pool several dozens of samples to one library, which extremely reduce per sample cost in a reasonable time. Thus, these next-generation sequencing based methods have been widely explored for the development of high-density linkage map in plants (Chen *et al.*, 2014; Chutimanitsakun *et al.*, 2011; Jia *et al.*, 2013; Pfender *et al.*, 2011; Saintenac *et al.*, 2013; Xie *et al.*, 2010; Zhang *et al.*, 2013).

Grain size or weight is genetically controlled by multiple genes, and large number of quantitative trait loci (QTL) for grain traits in wheat have been characterized in the past two decades (Brinton *et al.*, 2017; Gegas *et al.*, 2010; Kumar *et al.*, 2016; Maphosa *et al.*, 2014; Prashant *et al.*, 2012; Rasheed *et al.*, 2014; Tsilo *et al.*, 2010; Williams and Sorrells, 2014). The identified QTL distributed along all the wheat chromosomes, especially the stable and major QTL on A sub-genome 1A (Gegas *et al.*, 2010; Williams and Sorrells, 2014), 2A (Tsilo *et al.*, 2010), 3A (Gegas *et al.*, 2010; Kumar *et al.*, 2016), 4A (Prashant *et al.*, 2012), 5A (Brinton *et al.*, 2017; Gegas *et al.*, 2010; Williams and Sorrells, 2014), 6A (Gegas *et al.*, 2010; Maphosa *et al.*, 2014), and 7A (Kumar *et al.*, 2016; Tsilo *et al.*, 2010). Of those characterized QTL, one QTL for grain weight on chromosome 5A was further validated with two near-isogenic lines (NILs) and fine-mapped to a genetic interval of 4.32 cM corresponding to 74.6 Mb genomic sequences in Chinese Spring RefSeq v1.0 genome (Brinton *et al.*, 2017). However, it is a big challenge to underpin the causative genes in such a long genomic interval, due to functional redundancy (genetic buffering) of genes in three homoeologous genomes (A, B and D) and high repetitive nature of the wheat genome (International Wheat Genome Sequencing Consortium, 2014; Slade *et al.*, 2005). Up to now, most candidate genes for grain size and weight in wheat were characterized through homology-based cloning (Kumar *et al.*, 2016; Maphosa *et al.*, 2014). Several genes in rice have been proved to influence wheat grain size and weight, e.g. *GW2* (*TaGW2*), *GS3* (*TaGS-D1*), *CKX2* (*TaCKX6*), *GS5* (*TaGS5*), *TGW6* (*TaTGW6*), *GASR7* (*TaGASR7*) and *GIF1* (*TaCWI*) (Li and Yang, 2017). Apart from this, the starch and sucrose biosynthesis pathway unraveled in model species was proved to function in common wheat, e.g. *TaAGPL*, *TaAGPS* and *TaSus2* (Hou *et al.*, 2017; Jiang *et al.*, 2011). Thus, the reverse genetics is an efficient approach in wheat to characterize the underlying genetic components of morphogenesis (Li and Yang, 2017). Nevertheless, along with the rapid progress of genome sequencing, forward genetics would greatly facilitate the characterization of candidate genes involved in the development of grain size and weight of wheat.

Einkorn wheat, *Triticum monococcum* ssp. *monococcum* L. (A^m^A^m^, 2n = 2x = 14), the only cultivated diploid wheat, was domesticated from its wild species *T. monococcum* ssp. *boeoticum* (A^b^A^b^). As one of left untouched crops, the wild einkorn wheat grew in natural environment without human intensive selection for thousands of years (Jing *et al.*, 2007). It preserves a large number of phenotypic variations, which would facilitate the dissection of genetic architectures for agronomic important traits (International Wheat Genome Sequencing Consortium, 2014; Jing *et al.*, 2007; Zaharieva and Monneveux, 2014). The leaf rust resistant gene *Lr10*, the most important domestication gene *Q* and vernalization genes *Vrn1* and *Vrn2* were map-based cloned with the help of einkorn wheat (Feuillet *et al.*, 2003; Simons *et al.*, 2006; Stein *et al.*, 2000; Yan *et al.*, 2003; Yan *et al.*, 2004). Thus, the genome characteristics, highly polymorphism, and diversified traits make einkorn wheat a good model plant for gene discovery and breeding improvement in hexaploid wheat (*T. aestivum*, 2n = 6x = 42, AABBDD) (Shindo *et al.*, 2002; Stein *et al.*, 2000; Yan *et al.*, 2003).

To unravel the genetic architecture of grain traits in einkorn wheat, a recombinant inbred line (RIL) population of wild and cultivated einkorn wheat was explored to map QTL and characterize the underlying candidate genes. We exploited population SNPs through RAD-seq approach, developed a high-density genetic map, conducted a genome-wide QTL analysis for grain traits, and elucidated the candidate genes or gene pathways underlying the QTL based on comparative genomics and RNA sequencing (RNA-seq) analysis. The data revealed complex genetic components determining the grain size variation and the positive alleles retained across domestication in einkorn wheat. The whole-genome transcriptomic profiling further elucidated the candidate genes underlying QTL with significantly differential expressions between cultivated and wild einkorn wheat, and the superior alleles identified in this work provided opportunities for wheat genetic improvement.

## Materials and Methods

### Plant material and phenotyping

The 109 RIL population (F_10_) of *T. monococcum* ssp. *boeoticum* (KT1-1) × *T. monococcum ssp. monococcum* (KT3-5) were selected for linkage map construction and QTL mapping. This population and its parents were kindly provided by the KOMUGI Wheat Genetic Resources Databases of Japan.

### Field experiment and phenotyping

The RILs and their parents were grown with two replicates in a completely randomized block design at the experimental station of the Institute of Genetics and Developmental Biology, Chinese Academy of Sciences, Beijing (40°5’56”N and 116°25’8”E) in four successive years (2011, 2012, 2013, and 2014), and at the experimental station of Henan Agricultural University, Zhengzhou (34°51’52”N, 113°35’45”E) in one year (2014). These environments are designated E1, E2, E3, E4, and E5, respectively.

All RILs and their parents were planted in single 2-m rows with 40 cm between rows and 20 cm between individuals. The seeds were harvested from five randomly selected guarded plants from each line and replicate and then threshed by manual. Grain weight was determined using 100 grains with three replicates and transformed to thousand-grain weight (TGW, g). 100 grains for each RIL from each replicate were imaged and processed using a SC-G software (WSeen, Hangzhou, China), and the averages of the grain length (GL, mm), width (GW, mm), length / width (GLW), area (GA, mm^2^), and circumference (GC, mm) were calculated accordingly.

### RAD library construction and sequencing

Complexity-reduced genomic libraries prepared using the restriction endonuclease *Sbf*I (CCTGCAGG) has been reported in other species with large genomes (Chutimanitsakun *et al.*, 2011; Pfender *et al.*, 2011). The genomic DNA of RILs were sufficiently digested with *Sbf*I and processed into RAD libraries according to the protocol of Baird et al. (2008). We used a set of 16 barcoded adapters with sticky ends complimentary to 3’ overhang (TGCA-3’) created by *Sbf*I. The RAD libraries with an average size of 500 bp was constructed. For each library, 16 RILs were pooled together with each 6-bp barcode sequence to distinguish them and loaded to one lane, except for the seventh lane which contained 13 RILs and two parental lines. The RAD libraries were sequenced for single read (101 bp) on an Illumina Genome Analyzer IIx.

### RAD-seq data processing and SNP calling

RAD-seq data were processed by commands from Stacks v0.98 (Catchen *et al.*, 2011). Firstly, raw data were split into individuals according to first 6-bp barcode sequences of reads and filtered using sliding window methods by *process_radtags* with parameters of “–e sbfI -c -q –r –i fastq –E phred33” implemented in Stacks. If any read contained uncalled base or low phred33 quality scores less than 10 in any sliding window of 0.15 × read length were removed and discarded. Then, SNP calling from these tag sequences with *SbfI* site were carried out by ustacks, cstacks, and sstacks from the RIL population, Finally, the SNPs were transformed to genotypes, filtered with calling ratio > 40%, and applied for constructing linkage map.

### Genetic map construction

All the collected genotypes for the RILs were subjected to linkage mapping, and the distorted markers (Chi-square test *P* < 0.01, deviating from the expected 1:1 Mendelian segregation ratio) were excluded if these markers greatly affect the order of their neighbor markers or excessively change linkage distance. The linkage grouping and marker ordering were conducted using Joinmap V4.0 (Van Ooijen and Voorips, 2004) based on LOD ranging from 3.0 to 15.0 and MSTmap (Wu *et al.*, 2008), respectively. Recombination frequencies were converted into centiMorgan using Kosambi function (Kosambi, 1943). The final graphical linkage maps were generated using MapChart2.0 (Voorrips, 2002).

### Syntenic analysis

BLASTN was used to align the SNP markers against the physical maps of hexaploid wheat (IWGSC WGA v0.4; accessed from https://urgi.versailles.inra.fr/) and barley (IBSC RefSeqv1.0 pre-publication, http://webblast.ipk-gatersleben.de/barley_ibsc/downloads). We filtered BLASTN output data based on e-value < 1e^-10^ and query coverage ≥ 90%, respectively. Those hits that cannot meet the conditions were discarded. To balance the relationship bias of alignment in different genomes, different percentage identity thresholds were used, 91% for H genome of barley, 98%, 96%, and 96% for A, B, and D genome of wheat, respectively. The cleaned data of alignments were used to compare the genetic map with physical maps using OmicCircos (Hu *et al.*, 2014) implemented in R (R Core Team, 2016).

### Statistical analysis and QTL mapping

The broad-sense heritability (*H^2^*) of six quantitative traits was estimated with analysis of variance (ANOVA) and Pearson’s correlation coefficients among traits were calculated in R (R Core Team, 2016). The coefficient of variation (CV) values independently calculated for six traits from each individual environment as below: σ/μ, σ and μ represent the standard deviation and the mean of the phenotypic data in the population. QTL analysis were performed using composite interval mapping (CIM) method in Windows QTL Cartographer V2.5 (Wang *et al.*, 2012b) as previous procedure (Yu *et al.*, 2017). QTL were reported according to the previous method (Kumar *et al.*, 2016). Only the QTL detected above LOD threshold were included. If any significant QTL was identified with LOD below the threshold, but > 2 in other environments, the QTL were also included in the results as supporting information. Those QTL identified in at least two environments or associated with at least two traits were reported. QTL linked with flanking markers or overlapped confidence intervals (CIs, ± 1 LOD) were considered as one QTL for each trait with the CI reassigned by the overlapped genetic positions, while the unique genomic regions were considered as regions with at least one QTL included. The total explained variance by QTL were estimated using both ANOVA and multiple regression according to previous study (Yu *et al.*, 2017).

### RNA-seq and data analysis

#### RNA extraction and sequencing

At one-leaf stage seedlings from KT1-1 and KT3-5 were grown at 4∼6 °C treatment with six weeks. After that, the seedlings were transplanted to 25 cm^2^ × 25 cm pot in greenhouse under growing conditions of 16 h light and 25 °C and 8 h darkness and 15 °C. Spikes were harvested at 0, 7, 14, and 21 day(s) after flowering (DAF). From each developmental stage, more than three spikes from each line were pooled for RNA isolation. Each sample has three biological repeats. The construction of the libraries and RNA-seq were performed by the BGI (Shenzhen, China). The cDNA libraries with average insert size 300 bp from 24 samples were prepared with TruSeq RNA Sample Preparation Kit v2 (Illumina San Diego, USA) and sequenced on HiSeq4000 (Illumina, San Diego, USA) according to manufacturer’s standard protocols.

#### Differential gene expression analysis

The raw RNA-seq reads were filtered for contamination of the adaptor reads and low-quality reads or unknown nucleotides. The resultant clean reads were aligned against wheat accession Chinese Spring (CS) TGACv1 genome assembly (http://plants.ensembl.org/Triticum_aestivum). Transcript count information for sequences to each gene was calculated and normalized to the fragments per kilobase of transcript per million mapped reads (FPKM) values (Trapnell *et al.*, 2010). Significant differentially expressed (DE) genes were screened using bioconductor package NOISeq (Tarazona *et al.*, 2011). The genes with |log_2_(FPKM_KT3-5_/FPKM_KT1-1_)| > 1, and probability > 0.8 were identified as DE genes, while genes with the probability > 0.7 were considered as suggestive DE genes. Gene functions were assigned according to the best match of the alignments using BLASTP (v2.2.23) with default parameters to the Kyoto Encyclopedia of Genes and Genomes (KEGG) (http://www.genome.jp/kegg/), NR (ftp://ftp.ncbi.nlm.nih.gov/blast/db) and Gene Ontology (GO) databases. GO terms of the investigated genes were obtained by using Blast2GO (Conesa *et al.*, 2005), and KEGG pathways in which the genes might be involved were extracted from the matched genes in KEGG database. GO and KEGG pathway functional enrichment analysis were performed using phyper function implemented in R (R Core Team, 2016). GO terms with corrected-*P* values ≤ 0.05 and KEGG pathways with Q values ≤ 0.05 were defined as significant enrichments.

## Results

### Multigenic control of grain size related traits in einkorn wheat

To investigate the agronomical performance of grain size related traits, the 109 RILs derived from an inter sub-specific cross, *T. boeoticum* KT1-1 × *T. monococcum* KT3-5 (Shindo *et al.*, 2002; Yu *et al.*, 2017), were grown in five environments (designed from E1 to E5, respectively) under different agro-climatic conditions. The phenotypic data were collected, including grain length (GL), width (GW), length / width ratio (GLW), area (GA), circumference (GC), and thousand-grain weight (TGW) (**Table S1**; **Fig. 1**; **Fig. S1**). The wild einkorn wheat KT1-1 generally has small seed size, GL = 6.94 mm and GW = 1.79 mm in average, and the cultivated einkorn KT3-5 has big seed size, GL = 7.94 mm and GW = 2.64, with a 47% increase in GW than that of KT1-1. Among the RIL population, GL and GW distributed continuously and preserved a transgressive inheritance, for example, GL varied from 6.17 mm to 8.85 mm with a mean of 7.71 mm in E2. High broad-sense heritability (*H^2^*) was observed in both traits (0.86 for GL and 0.82 for GW) though phenotypic difference occurred among different environments. However, significant higher coefficient of variation (CV) were documented in GW across four environments (10.37% of GW versus 7.36% of GL, *P* = 0.0094 based on *t-test*), demonstrating that GW harbored larger variations in this einkorn wheat population. Moreover, TGW showed the highest CV from 21.16% (E1) to 32.64% (E3) across all environments, but with high heritability (0.85), revealing a relative larger effect of genotype-by-environment interactions. Meanwhile, the heritability of other traits were not lower than 0.80, though the phenotypic performance varied widely in different environments (**Table S1**). Thus, the large variations of the observed traits were proposed mainly under genetic control with multiple loci in this RIL population.

**Fig. 1.**
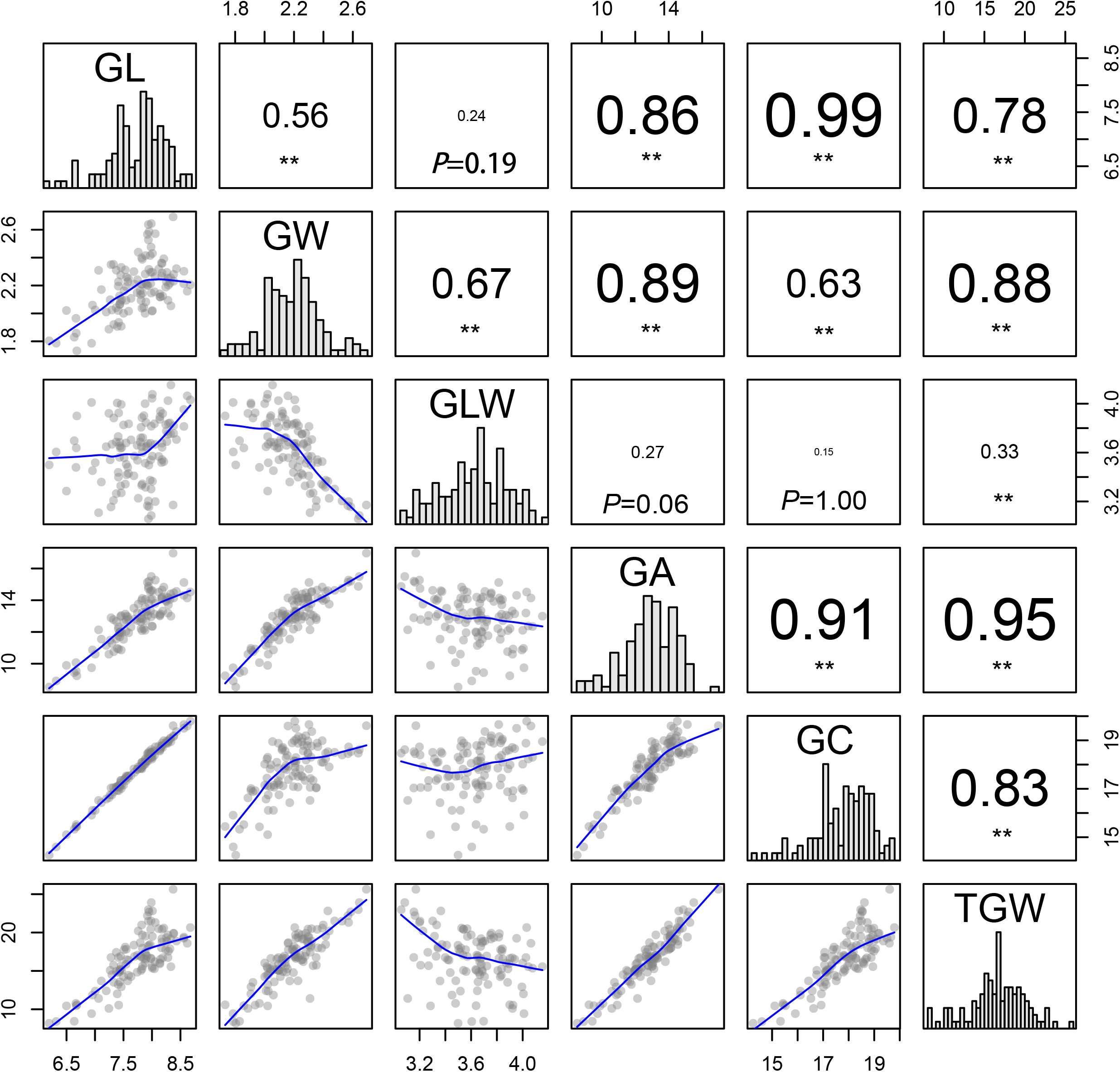
Phenotypic performances and distribution and correlation coefficients for six quantitative traits of parents and RILs using the average phenotypic data. The frequency distribution of phenotypic data of each trait was shown in the histogram at the diagonal cells. The X-Y scatter plot showed the correlation between traits at the lower-triangle panel, while the corresponding Pearson’s coefficients and *P* values of multiple comparison significant test were put on the upper-triangle panel. *, *P* < 0.05; **, *P* < 0.01. GL, grain length; GW, grain width; GLW, grain length / width; GA, grain area; GC, grain circumference; TGW, thousand-grain weight.

The significant correlations were observed among GL, GW, GLW, GA, GC and TGW (**Table S2**). GL had the highest positive correlation with GC (*r* = 0.99, Bonferroni-adjusted *P* < 0.01), followed by TGW versus GA (0.95, *P* < 0.01), GC versus GA (0.91, *P* < 0.01), and GW versus GA (0.89, *P* < 0.01) (**Fig. 1**). TGW positively correlated with all the grain size related traits, except GLW (−0.33, *P* < 0.01), of which a stronger correlation with GW was detected (from 0.73 to 0.93 with an average of 0.88, *P* < 0.01) than GL (from 0.66 to 0.82 with an average of 0.78, *P* < 0.01) across all the surveyed environments (**Fig. 1**; **Table S2**). Moreover, GW and GL had unbalanced correlations with the composite traits, GLW, GA and GC (**Fig. 1**). The GLW negatively correlated significantly with GW (−0.67, *P* < 0.01), but did not correlated with GL (0.24, *P* = 0.19). GC highly correlated with GL (0.99 (GL) *versus* 0.63 (GW), *P* < 0.01), while GA had an almost equal significant positive correlation with GL and GW. Thus, GW was a more important determinant of TGW in this RIL population, and GL and GW differentially contributed to these composite traits, GLW (grain shape), and GA and GC (grain size). This analysis revealed the complexity of genetic architectures and genetic connections of grain size related traits.

The principal component analysis (PCA) was performed to identify the major sources of variations in the morphometric datasets on the environment-wide dataset (**Fig. S2**; **Table S3**). Two extracted PCs, PC1 and PC2, significantly captured 97.0% to 98.6% of the total variations in the RIL population in each environment (**Table S3Error! Reference source not found.**). These two PCs showed similar organizations in four environments, of which PC1 (65.5% to 77.8%) and PC2 (20.8% to 31.5%) captured primary variation in grain size and grain shape, respectively (**Fig. S2**). In addition, both PCs with 72.7% (PC1) and 25.7% (PC2) of the explained variation were simultaneously identified on an environment-wide dataset (the average values and BLUP for each trait) (**Table S3**). Therefore, PC2 captures primarily grain shape differences including GLW, GW and GL, and PC1 describes grain size variations, where a proportional increase in GL and GW positively associates with an increase of GA and subsequently grain weight (**Fig. S2c**).

### Development of a high-density genetic map through RAD-sequencing (RAD-seq) approach

To construct a high-density genetic linkage map, RAD-seq platform was explored to characterize SNPs between two parents, and their RIL population. A *SbfI* reduced-representation library was constructed, and a total of 64.88 Gb sequences in seven lanes on Illumina Genome AnalyzerIIx were subsequently generated. These single-end reads (642 M) with 101 bp were demultiplexed to two parental lines and their 109 RILs according to the corresponding barcodes (**Table S4**). The reads with low base quality, or with ambiguous barcodes or *SbfI* cut-sites identified by *process_radtags* embedded in Stacks (Catchen *et al.*, 2011), were discarded. Finally, 438 M clean reads were trimmed to 94 bp per read after removing the first 6-bp barcode sequence and the last base, and this resulted ~3.95±1.06 M (mean ± standard deviance) reads per sample for further analysis (**Table S4**). The SNP calling using Stacks (Catchen *et al.*, 2011) with all the 438 M clean reads resulted in 42278 putative SNPs at 25805 genomic tags (11.36% of total 227244 tags). Of these SNPs, 25609 SNPs distributed at 16566 tags were retained with more than 40% calling rate in the RIL population, and these tags were hereinafter referred to as SNP markers for SNPs in one tag formed one haplotype.

These 16566 SNP markers were subjected to construct the genetic map, along with the available 939 markers (including DArT, SSR, Gene, and RFLP etc.) (Yu *et al.*, 2017). The resulted genetic map contained 10876 molecular markers distributed on seven chromosomes designated as Tm1A to Tm7A based on the known mapped markers, and these markers were grouped into 1551 unique bins (**Table 1**; **Fig. 2**). The genetic map spanned a total length of 1873.04 cM with an average marker interval of 0.17 cM and average marker density of 5.8 per cM (**Table 1**). The number of markers on each chromosome varied from 1,343 (Tm1A) and 1,732 (Tm2A) and genetic bins from 176 (Tm4A) to 265 (Tm7A) per chromosome. The shortest chromosome was Tm6A, and it harbored 1357 SNP and 115 other type markers with a genetic length of 238.6 cM, an average marker interval of 0.16 cM and marker density of 6.17 markers per cM. The longest chromosome was Tm3A, which contained 1656 markers with a genetic length of 293.5 cM, an average marker interval of 0.18 cM. On this map, the average bin length of each chromosome varied from 1.07 cM of Tm7A to 1.50 cM of Tm4A. Moreover, the map represented an average physical length of 454.21 kb (4.94 Gb/10876) when considering the genome length of *T. urartu* (Ling *et al.*, 2013).

**Fig. 2.**
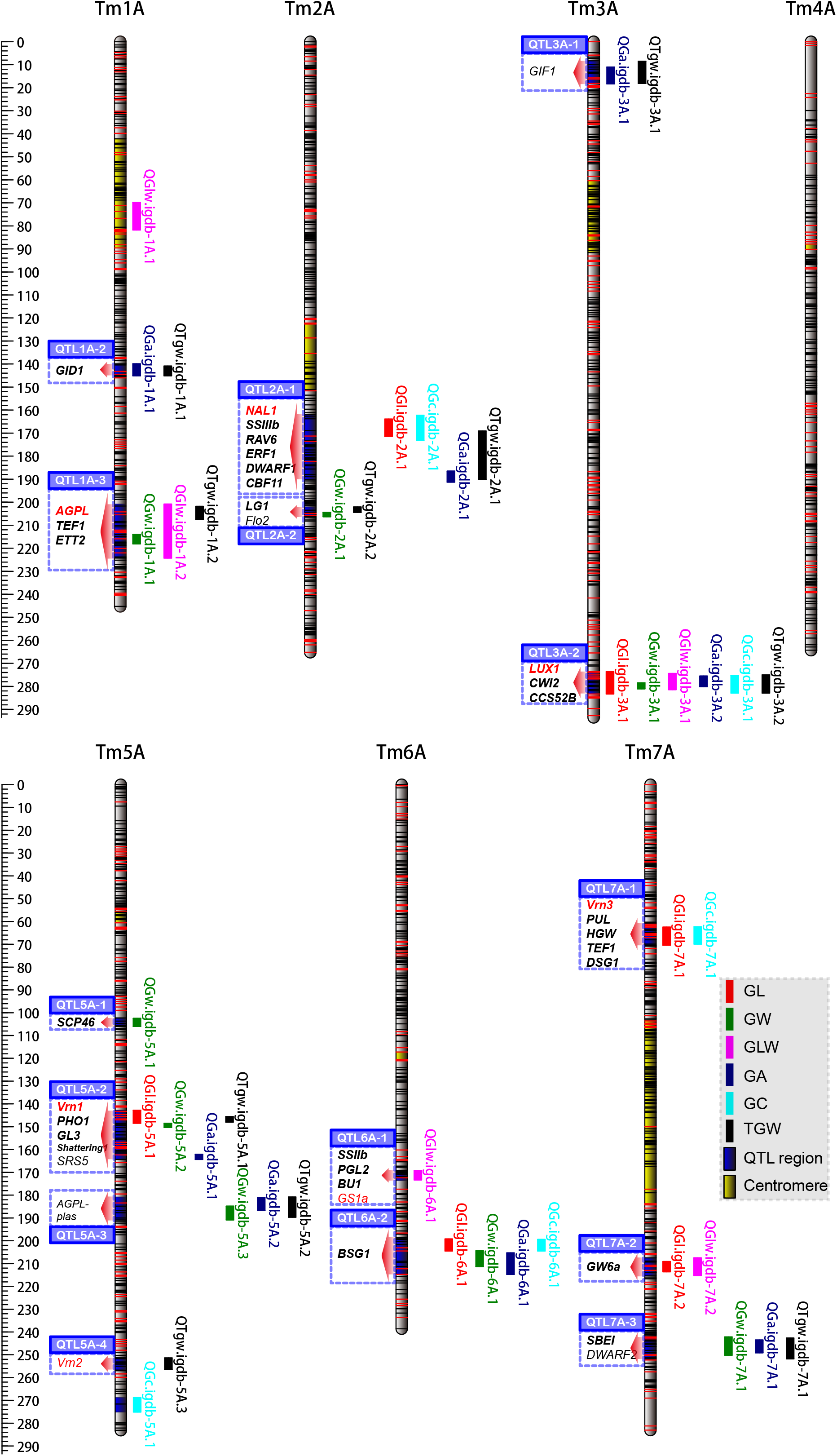
QTL detected in genome-wide using high-density genetic map of einkorn wheat. Genetic map showed 17 genomic regions harboring quantitative trait loci (QTL) for six grain traits in an einkorn wheat RIL population of *T. monococcum* ssp. *boeoticum* (KT1-1) and *T. monococcum* ssp. *monococcum* (KT3-5). The detected QTL for each trait from each environment were combined with confidential intervals and mapped on the genetic map. At each linkage group, each QTL was plotted at right side, while the candidate genes in each QTL region were put at left side. Detailed information of QTL is available in **Error! Reference source not found.**. The candidate genes of each QTL region were shown in blue shaded, the red arrows showing the QTL regions. Genes in red were mapped through developing functional markers and genetic mapping, while black bold denoted genes located inside QTL region, red or black normal for genes surrounding QTL region. The yellow shaded portions of each linkage group are the probable centromere regions. The positions of SNP and other type of markers were denoted with black and red ticks, respectively. GL, grain length, GW, grain width, GLW, grain length / width, GA, grain area, GC, grain circumference, TGW, thousand-grain weight.

**Table 1.** Summary information of high-density of the high-density einkorn wheat genetic map.

| Chr. | Length (cM) | SNP | Other<br>markers | Total marker<br>No. | Bin No. | Bin length<br>(cM) | Marker interval<br>(cM) | Max. gap<br>(cM) |
| --- | --- | --- | --- | --- | --- | --- | --- | --- |
| Tm1A | 245.33 | 1,233 | 110 | 1,343 | 206 | 1.19 | 0.18 | 4.85 |
| Tm2A | 265.16 | 1,621 | 111 | 1,732 | 230 | 1.15 | 0.15 | 6.71 |
| Tm3A | 293.50 | 1,466 | 190 | 1,656 | 242 | 1.21 | 0.18 | 6.28 |
| Tm4A | 264.33 | 1,453 | 103 | 1,556 | 176 | 1.50 | 0.17 | 20.82 |
| Tm5A | 283.08 | 1,269 | 153 | 1,422 | 245 | 1.16 | 0.20 | 8.16 |
| Tm6A | 238.61 | 1,357 | 115 | 1,472 | 187 | 1.28 | 0.16 | 5.02 |
| Tm7A | 283.04 | 1,538 | 157 | 1,695 | 265 | 1.07 | 0.17 | 12.17 |
| Total | 1,873.04 | 9,937 | 939 | 10,876 | 1,551 | 1.21 | 0.17 | 20.82 |

Compared with previous map constructed with the same RIL population (Yu *et al.*, 2017), this high-density SNP genetic map has been extended about 496 cM (from 1377 cM to 1873 cM) through mapping SNP markers in-between other markers or on the distal ends of linkage groups. The length extension was primarily resulted from additional intra-chromosomal recombination (~439 cM) detected by the new mapped SNP markers. Meanwhile, 84 SNP markers located beyond other markers at the distal ends of chromosomes, of which twenty-one were mapped on Tm1AS/L, six on Tm2AS, two on Tm4AL, fifty-one on Tm5AS/L, and four on Tm6AS. These extensions covered 29 bins with a length of ~57 cM in the SNP linkage map. Regarding to the gap in this high-density linkage map, all the intervals had a length of <10 cM between two neighboring markers, albeit one gap on Tm4A (20.82 cM between the bin3 to bin4) and two gaps on Tm7A (12.17 cM between bin263 and bin264, 10.02 cM between bin173 and bin174) (**Fig. 2**; **Table S5**). We noticed only one gap greater than 20 cM on Tm4A, while the previous reported two gaps (> 17 cM) were saturated with SNP markers. The gap on Tm4A should be the properties of this RIL population (Hori *et al.*, 2007; Shindo *et al.*, 2002; Yu *et al.*, 2017), and even shared by other einkorn wheat population (Jing *et al.*, 2007; Singh *et al.*, 2007). Thus, this einkorn wheat genetic linkage map was greatly improved in marker density and evenness and represented to be a high-quality map with thousands of markers and limited gaps on each linkage group, which provided an elite tool for unraveling the genetic components of agronomic important traits and genome assembling of einkorn wheat.

### Homologous regions in barley and wheat

To evaluate the quality and genome coverage of our SNP genetic map, 9937 SNP markers were aligned via BLASTN against barley (IBSC RefSeqv1.0, referred as HvRefSeqV1) (Mascher *et al.*, 2017) and hexaploid wheat genomes (accession “Chinese Spring”, IWGSC WGA v0.4). The parameter of identities, 98% for A genome, 96% for B and D genomes, and 91% for H genome were used to filter alignments to decrease the marker number bias because of the relatedness of these four genomes. This resulted in 1834, 2187, 1693, 922 hits at A, B, D and H genomes, respectively. The filtered alignments corresponded to 3102 SNP markers, 88.91% of which were syntenous and mapped to the expected homologous chromosomes. Through Spearman’s rank correlation coefficient (ρ) of the 2758 syntenous marker locations on einkorn wheat and four genomes, the high level of collinearity (average ρ = 0.76) between physical maps and genetic distances of einkorn wheat was verified, except Tm4A *vs* Ta4A and Tm7A *vs* Ta7B (**Fig. 3a, b**). This should be attributed to 4AL/5AL/7BS translocations during wheat’s evolutionary history (Devos *et al.*, 1995; King *et al.*, 1994). The translocation between 4A and 5A was observed in einkorn wheat when compared the genetic map with B, D, and H genomes (**Fig. 3b**; **Fig. S3a, b**). This translocation corresponded to ~50 cM in the genetic map and ~35 Mb, ~28 Mb, ~20 Mb in the chromosomes Ta4B, Ta4D, and Hv4H, respectively (**Fig. 3c**). However, the 4AL/7BS translocation were not obviously detected using our data for only two SNPs from the distal end of Tm4A covering ~1 cM were mapped onto Ta7BS (**Fig. 3c**). Moreover, the gene-rich regions preserved higher collinearity than centromeric regions, where the shortness of genetic distance was shown because of less recombination (**Fig. 3a**). It was worth noting that the mapped SNP markers covered > 97% of all the genomes, except for Ta4A (92%) and Ta3B (93%), revealing a high genome coverage of this genetic map (**Fig. S3c**). Therefore, the high collinearity provided the syntenic blocks between einkorn wheat and the available genomes, which would facilitate the identification of the interesting genomic blocks for further analysis.

**Fig. 3.**
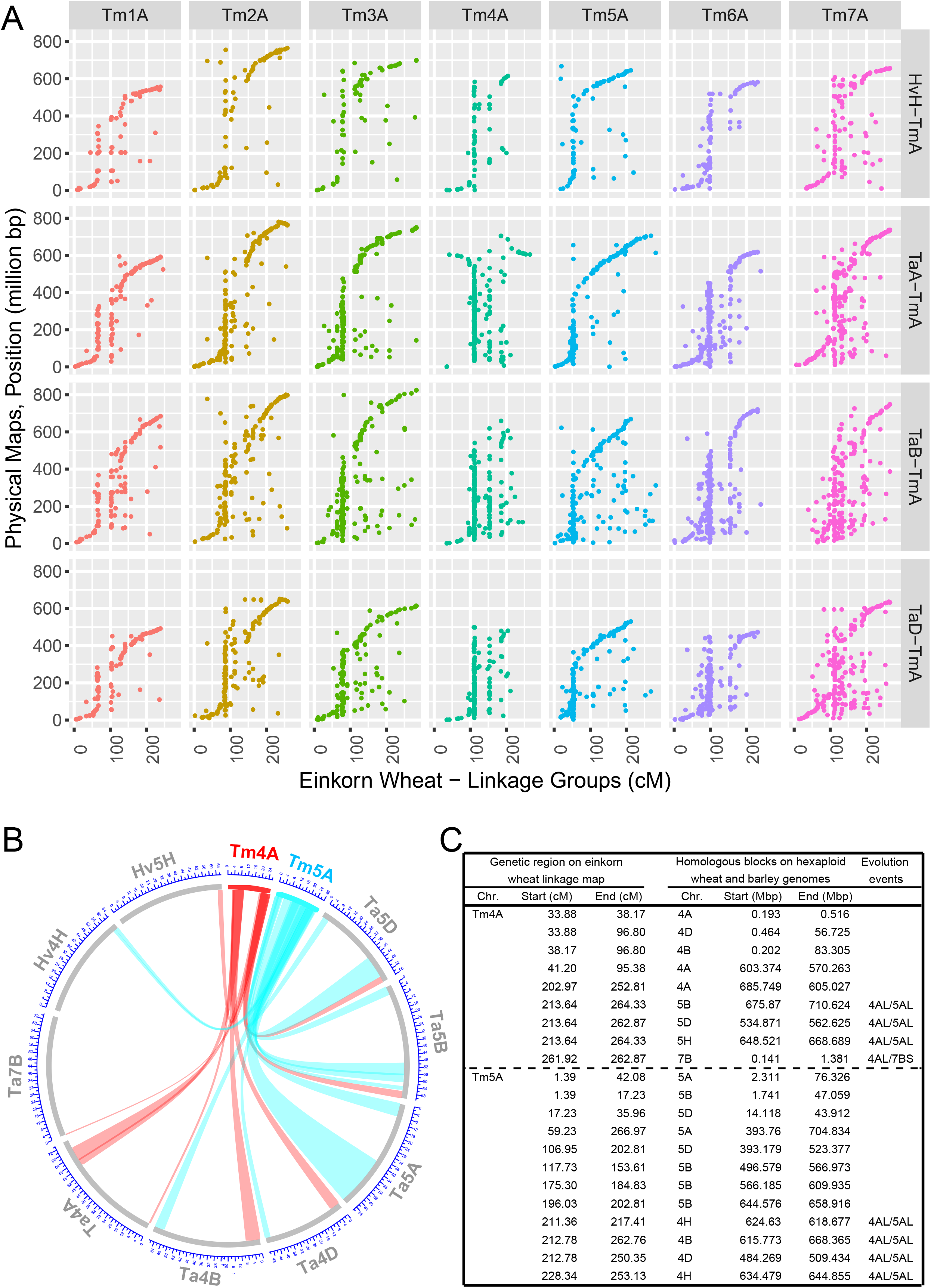
Genomic collinearity and chromosomal structure variations revealed by using the high-density genetic map of einkorn wheat. (a) SNP markers were aligned against the four einkorn wheat related genomes (A, B, D from hexaploid wheat, and H genome from barley), and the positions of the hit markers were compared with physical locations from four genomes. (b) Comparisons of the marker positions on homologous groups 4, 5, and 7, elucidate 4AL/5AL/7BS translocations using einkorn wheat genetic map. The detail information were given in (c).

### Genetic architectures of grain size related traits in einkorn wheat

To elucidate the genetic architecture of grain size related traits in einkorn wheat, a genome-wide QTL analysis through Windows QTL Cartographer (Wang *et al.*, 2012b) was performed in this RIL population, along with the phenotypic data and the high-density SNP linkage map. Using the CIM method, a total of 42 additive QTL were identified in five environments distributed across six chromosomes, except Tm4A, and they had a LOD peak score of 3.4 or more and explained 6.4% to 38.1% of the phenotypic variations (**Fig. S4**; **Table S6**). Among 42 QTL, 31 (74%) loci involved alleles from KT3-5 for increasing phenotypic values, while the other 11 (26%) loci had alleles from KT1-1 for increasing phenotypic values, suggested that positive alleles for grain size related traits were present even in the parent with low phenotypic value. These 42 unique QTL were assigned to 17 genomic regions for some QTL for several traits co-located at the same region on chromosomes. This resulted an average number of 2.5 QTL for each region, of which 3A-2 (273.6-282.9 cM) harbored the most of six traits related QTL (**Table 2**; **Fig. 2**).

**Table 2.**
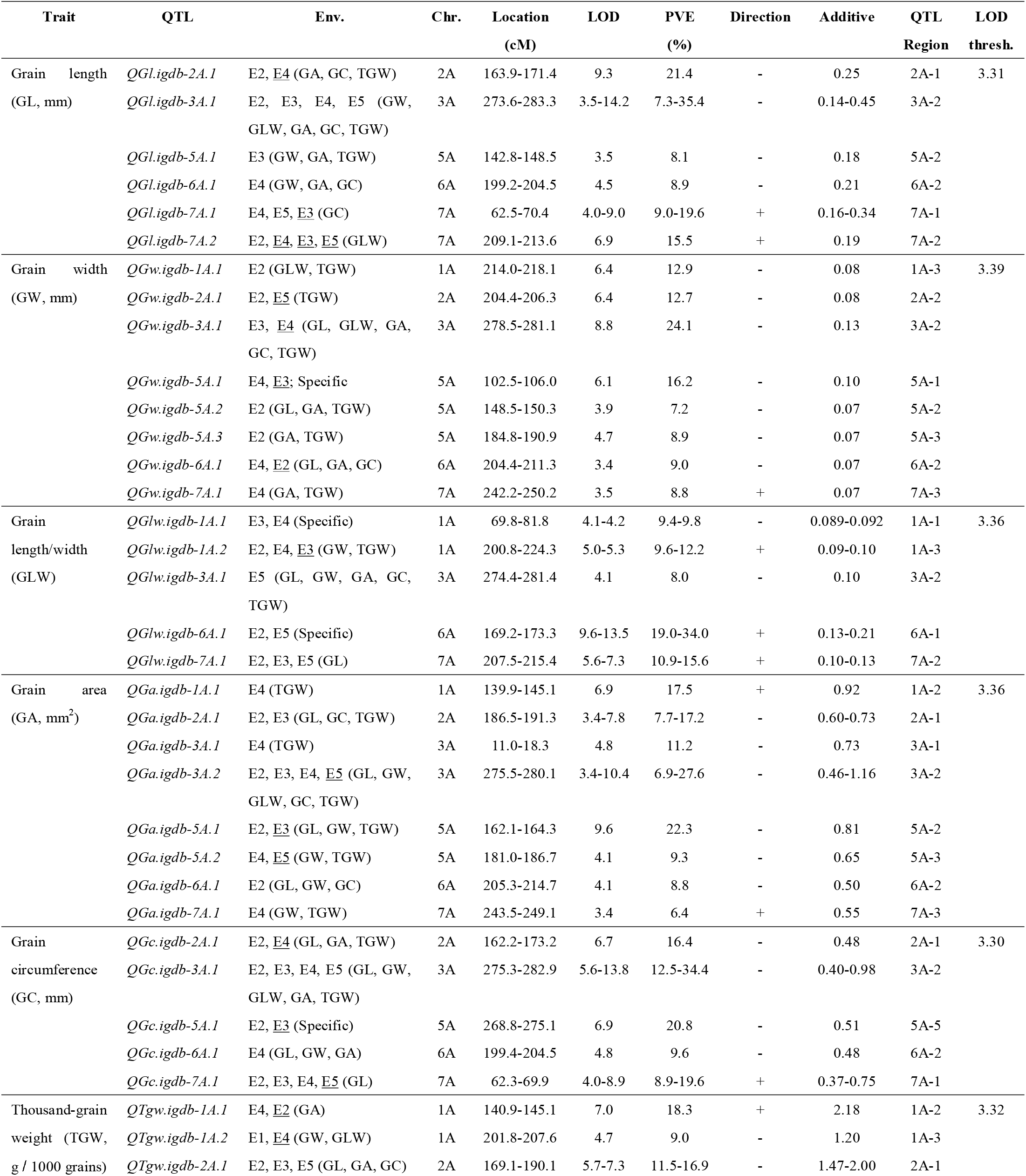
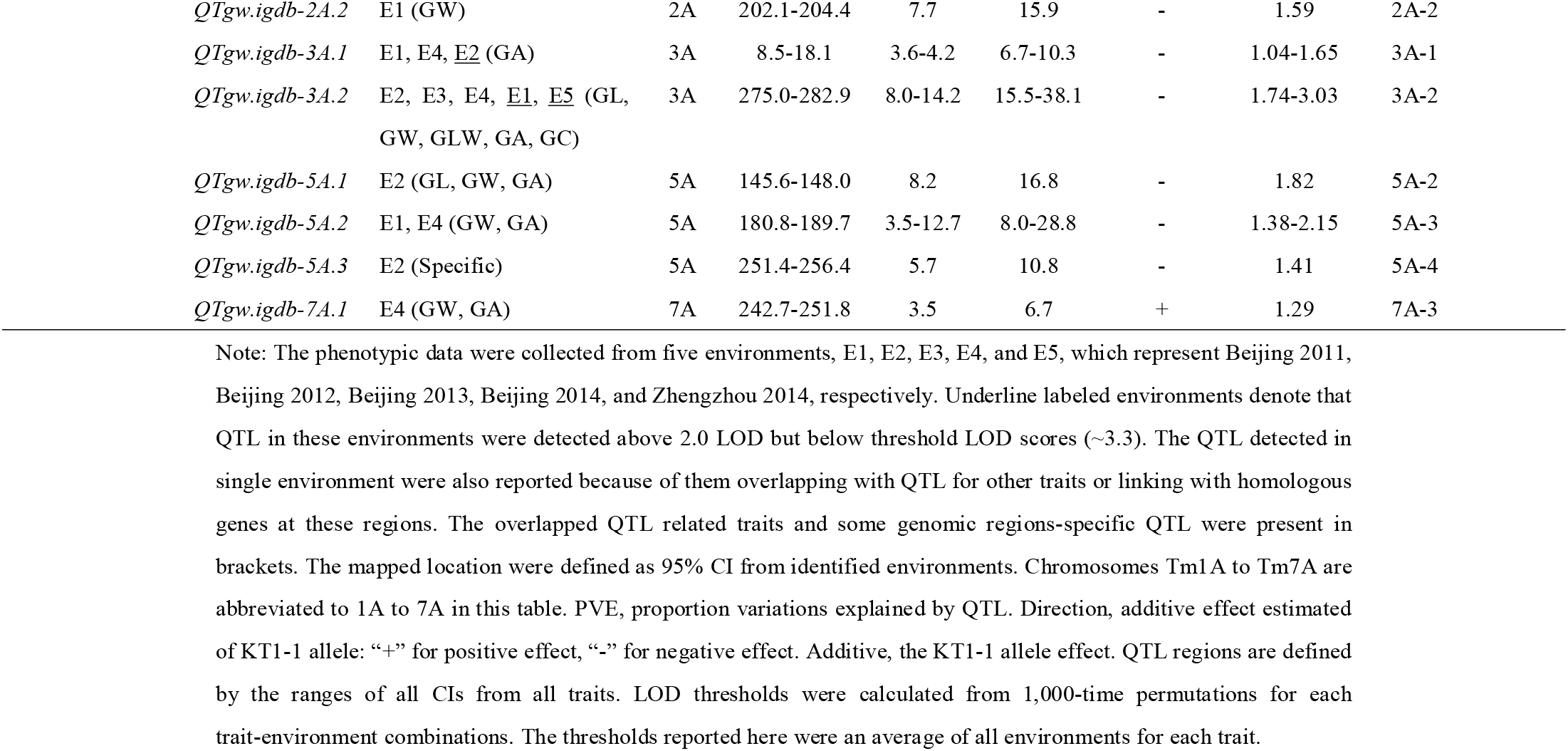
QTL detected with CIM method using the high-density einkorn wheat genetic map.

| Trait | QTL | Env. | Chr. | Location<br>(cM) | LOD | PVE<br>(%) | Direction | Additive | QTL<br>Region | LOD<br>thresh. |
| --- | --- | --- | --- | --- | --- | --- | --- | --- | --- | --- |
| Grain length<br>(GL, mm) | *QGl.igdb-2A.1* | E2, <u>E4</u> (GA, GC, TGW) | 2A | 163.9-171.4 | 9.3 | 21.4 | - | 0.25 | 2A-1 | 3.31 |
|  | *QGl.igdb-3A.1* | E2, E3, E4, E5 (GW, GLW, GA, GC, TGW) | 3A | 273.6-283.3 | 3.5-14.2 | 7.3-35.4 | - | 0.14-0.45 | 3A-2 |  |
|  | *QGl.igdb-5A.1* | E3 (GW, GA, TGW) | 5A | 142.8-148.5 | 3.5 | 8.1 | - | 0.18 | 5A-2 |  |
|  | *QGl.igdb-6A.1* | E4 (GW, GA, GC) | 6A | 199.2-204.5 | 4.5 | 8.9 | - | 0.21 | 6A-2 |  |
|  | *QGl.igdb-7A.1* | E4, E5, <u>E3</u> (GC) | 7A | 62.5-70.4 | 4.0-9.0 | 9.0-19.6 | + | 0.16-0.34 | 7A-1 |  |
|  | *QGl.igdb-7A.2* | E2, <u>E4</u>, <u>E3</u>, <u>E5</u> (GLW) | 7A | 209.1-213.6 | 6.9 | 15.5 | + | 0.19 | 7A-2 |  |
| Grain width<br>(GW, mm) | *QGw.igdb-1A.1* | E2 (GLW, TGW) | 1A | 214.0-218.1 | 6.4 | 12.9 | - | 0.08 | 1A-3 | 3.39 |
|  | *QGw.igdb-2A.1* | E2, <u>E5</u> (TGW) | 2A | 204.4-206.3 | 6.4 | 12.7 | - | 0.08 | 2A-2 |  |
|  | *QGw.igdb-3A.1* | E3, <u>E4</u> (GL, GLW, GA, GC, TGW) | 3A | 278.5-281.1 | 8.8 | 24.1 | - | 0.13 | 3A-2 |  |
|  | *QGw.igdb-5A.1* | E4, <u>E3</u>; Specific | 5A | 102.5-106.0 | 6.1 | 16.2 | - | 0.10 | 5A-1 |  |
|  | *QGw.igdb-5A.2* | E2 (GL, GA, TGW) | 5A | 148.5-150.3 | 3.9 | 7.2 | - | 0.07 | 5A-2 |  |
|  | *QGw.igdb-5A.3* | E2 (GA, TGW) | 5A | 184.8-190.9 | 4.7 | 8.9 | - | 0.07 | 5A-3 |  |
|  | *QGw.igdb-6A.1* | E4, <u>E2</u> (GL, GA, GC) | 6A | 204.4-211.3 | 3.4 | 9.0 | - | 0.07 | 6A-2 |  |
|  | *QGw.igdb-7A.1* | E4 (GA, TGW) | 7A | 242.2-250.2 | 3.5 | 8.8 | + | 0.07 | 7A-3 |  |
| Grain length/width<br>(GLW) | *QGlw.igdb-1A.1* | E3, E4 (Specific) | 1A | 69.8-81.8 | 4.1-4.2 | 9.4-9.8 | - | 0.089-0.092 | 1A-1 | 3.36 |
|  | *QGlw.igdb-1A.2* | E2, E4, <u>E3</u> (GW, TGW) | 1A | 200.8-224.3 | 5.0-5.3 | 9.6-12.2 | + | 0.09-0.10 | 1A-3 |  |
|  | *QGlw.igdb-3A.1* | E5 (GL, GW, GA, GC, TGW) | 3A | 274.4-281.4 | 4.1 | 8.0 | - | 0.10 | 3A-2 |  |
|  | *QGlw.igdb-6A.1* | E2, E5 (Specific) | 6A | 169.2-173.3 | 9.6-13.5 | 19.0-34.0 | + | 0.13-0.21 | 6A-1 |  |
|  | *QGlw.igdb-7A.1* | E2, E3, E5 (GL) | 7A | 207.5-215.4 | 5.6-7.3 | 10.9-15.6 | + | 0.10-0.13 | 7A-2 |  |
| Grain area<br>(GA, mm<sup>2</sup>) | *QGa.igdb-1A.1* | E4 (TGW) | 1A | 139.9-145.1 | 6.9 | 17.5 | + | 0.92 | 1A-2 | 3.36 |
|  | *QGa.igdb-2A.1* | E2, E3 (GL, GC, TGW) | 2A | 186.5-191.3 | 3.4-7.8 | 7.7-17.2 | - | 0.60-0.73 | 2A-1 |  |
|  | *QGa.igdb-3A.1* | E4 (TGW) | 3A | 11.0-18.3 | 4.8 | 11.2 | - | 0.73 | 3A-1 |  |
|  | *QGa.igdb-3A.2* | E2, E3, E4, <u>E5</u> (GL, GW, GLW, GC, TGW) | 3A | 275.5-280.1 | 3.4-10.4 | 6.9-27.6 | - | 0.46-1.16 | 3A-2 |  |
|  | *QGa.igdb-5A.1* | E2, <u>E3</u> (GL, GW, TGW) | 5A | 162.1-164.3 | 9.6 | 22.3 | - | 0.81 | 5A-2 |  |
|  | *QGa.igdb-5A.2* | E4, <u>E5</u> (GW, TGW) | 5A | 181.0-186.7 | 4.1 | 9.3 | - | 0.65 | 5A-3 |  |
|  | *QGa.igdb-6A.1* | E2 (GL, GW, GC) | 6A | 205.3-214.7 | 4.1 | 8.8 | - | 0.50 | 6A-2 |  |
|  | *QGa.igdb-7A.1* | E4 (GW, TGW) | 7A | 243.5-249.1 | 3.4 | 6.4 | + | 0.55 | 7A-3 |  |
| Grain circumference<br>(GC, mm) | *QGc.igdb-2A.1* | E2, <u>E4</u> (GL, GA, TGW) | 2A | 162.2-173.2 | 6.7 | 16.4 | - | 0.48 | 2A-1 | 3.30 |
|  | *QGc.igdb-3A.1* | E2, E3, E4, E5 (GL, GW, GLW, GA, TGW) | 3A | 275.3-282.9 | 5.6-13.8 | 12.5-34.4 | - | 0.40-0.98 | 3A-2 |  |
|  | *QGc.igdb-5A.1* | E2, <u>E3</u> (Specific) | 5A | 268.8-275.1 | 6.9 | 20.8 | - | 0.51 | 5A-5 |  |
|  | *QGc.igdb-6A.1* | E4 (GL, GW, GA) | 6A | 199.4-204.5 | 4.8 | 9.6 | - | 0.48 | 6A-2 |  |
|  | *QGc.igdb-7A.1* | E2, E3, E4, <u>E5</u> (GL) | 7A | 62.3-69.9 | 4.0-8.9 | 8.9-19.6 | + | 0.37-0.75 | 7A-1 |  |
| Thousand-grain weight<br>(TGW, g / 1000 grains) | *QTgw.igdb-1A.1* | E4, <u>E2</u> (GA) | 1A | 140.9-145.1 | 7.0 | 18.3 | + | 2.18 | 1A-2 | 3.32 |
|  | *QTgw.igdb-1A.2* | E1, <u>E4</u> (GW, GLW) | 1A | 201.8-207.6 | 4.7 | 9.0 | - | 1.20 | 1A-3 |  |
|  | *QTgw.igdb-2A.1* | E2, E3, E5 (GL, GA, GC) | 2A | 169.1-190.1 | 5.7-7.3 | 11.5-16.9 | - | 1.47-2.00 | 2A-1 |  |

|  |  |  |  |  |  |  |  |  |
| --- | --- | --- | --- | --- | --- | --- | --- | --- |
| *QTgw.igdb-2A.2* | E1 (GW) | 2A | 202.1-204.4 | 7.7 | 15.9 | - | 1.59 | 2A-2 |
| *QTgw.igdb-3A.1* | E1, E4, <u>E2</u> (GA) | 3A | 8.5-18.1 | 3.6-4.2 | 6.7-10.3 | - | 1.04-1.65 | 3A-1 |
| *QTgw.igdb-3A.2* | E2, E3, E4, <u>E1</u>, <u>E5</u> (GL, GW, GLW, GA, GC) | 3A | 275.0-282.9 | 8.0-14.2 | 15.5-38.1 | - | 1.74-3.03 | 3A-2 |
| *QTgw.igdb-5A.1* | E2 (GL, GW, GA) | 5A | 145.6-148.0 | 8.2 | 16.8 | - | 1.82 | 5A-2 |
| *QTgw.igdb-5A.2* | E1, E4 (GW, GA) | 5A | 180.8-189.7 | 3.5-12.7 | 8.0-28.8 | - | 1.38-2.15 | 5A-3 |
| *QTgw.igdb-5A.3* | E2 (Specific) | 5A | 251.4-256.4 | 5.7 | 10.8 | - | 1.41 | 5A-4 |
| *QTgw.igdb-7A.1* | E4 (GW, GA) | 7A | 242.7-251.8 | 3.5 | 6.7 | + | 1.29 | 7A-3 |

Note: The phenotypic data were collected from five environments, E1, E2, E3, E4, and E5, which represent Beijing 2011, Beijing 2012, Beijing 2013, Beijing 2014, and Zhengzhou 2014, respectively. Underline labeled environments denote that QTL in these environments were detected above 2.0 LOD but below threshold LOD scores (~3.3). The QTL detected in single environment were also reported because of them overlapping with QTL for other traits or linking with homologous genes at these regions. The overlapped QTL related traits and some genomic regions-specific QTL were present in brackets. The mapped location were defined as 95% CI from identified environments. Chromosomes Tm1A to Tm7A are abbreviated to 1A to 7A in this table. PVE, proportion variations explained by QTL. Direction, additive effect estimated of KT1-1 allele: “+” for positive effect, “-” for negative effect. Additive, the KT1-1 allele effect. QTL regions are defined by the ranges of all CIs from all traits. LOD thresholds were calculated from 1,000-time permutations for each trait-environment combinations. The thresholds reported here were an average of all environments for each trait.

To investigate the candidate genes underlying these QTL, homologous genes with functions on grain size or weight reported in rice, barley, and wheat, were retrieved and mapped to our high-density genetic map. Except 1A-1 that is homologous to chromosomal centromeric region, the remaining sixteen QTL regions in einkorn wheat had homologous blocks in hexaploid wheat (Chinese Spring) and barley genomes. These syntenic blocks had average physical lengths of 24.29 Mb and 18.85 Mb in hexaploid wheat A and barley H genomes, respectively (**Fig. S5**). This process allowed 41 collected genes to be mapped, of which 40 and 36 homologs were detected in wheat and barley genomes, respectively (**Table S7**). Among all these genes, 30 genes were mapped in the QTL confidence intervals (CIs), seven genes (*Flo2*, *GIF1*, *SRS5*, *AGPL-plas*, *Vrn2*, *GS1a*, and *DWARF2*) were closely linked with their target QTL, but four genes had genetic distance larger than 10 cM from the identified QTL. These four genes corresponded to five genomic loci, *Sus2* (83.67 cM) and *GW7* (75.70 cM) on Tm2A, *Sus1* (32.65 cM), *Sus2* (83.67 cM), and *GASR7* (103.17 cM) on Tm7A (**Fig. 2**; **Table S7**). To confirm their concordant locations on the map, *AGPL*, *Sus1*, *Sus2*, *Vrn1*, *Vrn2*, *Vrn3*, *NAL1*, *GS1a*, *GASR7*, and *GW7* genes were mapped by developing polymorphic markers according to sequence variations (ie. SNP and InDel) (**Table S8**). One InDel was identified at the promotor region of *Vrn3*, and the resultant InDel marker was mapped to the homologous region on Tm7A as expected, closely linked with *PUL*, *HGW*, *TEF1* and *DSG1* genes (**Fig. 2**; **Fig. S6**). Collectively, *AGPL*, *Vrn1*, *Vrn3*, and *NAL1* were mapped to QTL regions 1A-3, 5A-2, 7A-1, and 2A-1, respectively, and *Vrn2* and *GS1a* were located to a small genetic distance of < 5 cM with 5A-2 and 6A-1, consistent with the comparative homologous regions. Thus, the genetic mapping of these genes further confirmed the homology-based mapping data and their sequence variations might affect their functions on grain size development.

#### GL & GW QTL

Cultivated einkorn wheat KT3-5 had longer and much wider grain than KT1-1, the other parent of the RIL population (**Table S1**). In total, eleven genomic regions were mapped QTL for both GL and GW (**Fig. S7**). For GL, six unique QTL were distributed over chromosomes Tm2A, Tm3A, Tm5A, Tm6A, and Tm7A, explaining from 7.27% to 35.43% of phenotypic variation across all environments (**Table S6**). The KT1-1 alleles on Tm2A, Tm3A, Tm5A and Tm6A decreased GL with 0.14-0.45 mm, while the allele on Tm7A increased 0.16-0.34 mm. All the detected QTL represented by peak markers could explain 59.1% of total phenotypic variation (**Table S9**). Multiple comparison of phenotypic data showed that when RILs inherited with 2-3 positive alleles, it would increase GL significantly with *P* < 0.01 (**Fig. S8**). Eight QTL for GW were detected across all six QTL-located chromosomes, jointly explaining 56.8% of total phenotypic variation of GW (**Table S9**). QTL of GW showed negative effect of most KT1-1 alleles decreasing grain width with 0.07-0.13 mm, and only *QGw.igdb-7A.1* mapped in 242.2-250.2 cM of 7A-3 showed increasing grain width of 0.07 mm (**Table 2**). Three QTL regions containing QTL for both GL and GW, were 3A-2, 5A-2, and 6A-2, and these regions located multiple genes including *TmLUX1*, *CWI2*, *CCS52B* for 3A-2, *Vrn1*, *PHO1*, *Shattering1*, *GL3* on 5A-2, *BSG1* on 6A-2 (**Fig. 2**). For GL, 2A-1 contained *NAL1*, 7A-1 for several genes including *Vrn3*, *PUL*, *HGW*, *TEF1*, *DSG1*, and 7A-2 for *GW6a*. For GW, 1A-3 had *AGPL*, *TEF1*, *ETT*, 7A-3 for *SBEI*, and the remaining QTL overlapped with QTL for other traits. Of these genes, *Vrn1*, *Vrn3*, *AGPL1*, *TmLUX1* and *NAL1*, were genetic mapped as previously reported (Yu *et al.*, 2017). These mapped genes showed large sequence variations between the two parental lines, indicating that they might be the candidate genes underlying the QTL and providing potential diagnostic markers for allelic selection. A 9-bp deletion in *AGPL1* promoter region (~1 kb upstream) was observed in KT1-1, and a SNP changed amino acid from S in KT3-5 to G in KT1-1, but other SNPs detected in other five exons were synonymous mutations (**Fig. S9**). Moreover, the increase of positive alleles of the detected nine QTL for GW showed more significant divergence between different groups than alleles for GL (**Fig. S8**; **Table S10**).

#### GLW, GA & GC QTL

The composite traits, GLW, GA, and GC, were calculated according to GL and GW and also exploited for QTL analysis, as they directly reflected grain size (GC, GA) and shape (GLW). Out of QTL for GLW, the most significant QTL *QGlw.igdb-6A.1*, can explain 19.0%-34.0% of the phenotypic variations (**Table 2**). This QTL was specifically mapped to genomic region 6A-1 that contains homologous genes *SSIIb*, *PGL2*, and *BU1*, which participated in starch biosynthesis and *brassinosteroid signaling*, and associated with starch accumulation and grain length and weight. The unique QTL region for GLW was 1A-1, which was generally syntenic with a large proportion of centromeric region of the physical maps and little information of homologous genes was available. For GA, *QGa.igdb-1A.1* located on 139.9-145.1 cM of Tm1A and explained 17.54% of the total phenotypic variation with positive effect with KT1-1 allele, which contained no QTL for GL and GW (**Table 2**). However, this QTL overlapped with TGW QTL *QTgw.igdb-1A.1*, and in this region, one gene for GID1 that interacted with RHT1 in plants to control the plant height was located. Moreover, the *QGa.igdb-3A.1* overlapped with *QTgw.igdb-3A.1* on 3A-1 that linked with *GIF1* (**Fig. 2**). For GC, four overlapped with GL QTL, consistent with strong correlation between these two traits (*r* = 0.99, *P* < 0.01). Overall, genetic overlaps were observed between the three composite traits and GL/GW, which were revealed by co-location of QTL for GL/GW with QTL for GLW (three of five), GA (six of eight), and GC (four of five).

#### TGW QTL

To understand the influence of grain size on grain weight, QTL analysis for TGW was conducted and ten QTL were identified on Tm1A, Tm2A, Tm3A, Tm5A, and Tm7A (**Table 2**). Two significant QTL, *QTgw.igdb-1A.1* and *QTgw.igdb-7A.1*, showed positive effects of KT1-1 allele on TGW with increasing thousand-grain weight of 2.18 g and 1.29 g, respectively (**Table 2**). The remaining QTL decreased the TGW of 1.04-3.03 g and explained 6.71% to 38.05% of the total phenotypic variations. Several previous mapped genes had effects on TGW or its related traits, like *Vrn1* (5A-2) and *Vrn3* (7A-1), which acted on opposite additive effects (**Table 2**). Although one QTL, *QTgw.igdb-5A.3* were mapped between 251.4-256.4 cM, less information was known according to homologous analysis, except that *Vrn2* (248.03 cM) closely linked with this region (**Table 2**). Interestingly, genes in the starch biosynthesis pathway, *AGPL*, *AGPL-plas*, *SSIIIb*, *PHO1* and *SBEI* being mapped five genomic regions 1A-3, 5A-3, 2A-1, 5A-2 and 7A-3, respectively, had negative effects of wild einkorn wheat KT1-1 allele, except 7A-3 for positive effect (**Fig. 2**). With respect to five grain size related traits, nine of TGW QTL overlapped, out of which seven of eight GA QTL and six of eight GW QTL coincided while only a half of GL QTL and two GW-associated GLW QTL were co-located. Along with correlation analysis and PCA of TGW and other traits, it further demonstrated that TGW was a complex trait and mainly determined by grain size. The data also demonstrated that grain size related traits (especially for GA and GW), associated positively more with the TGW, while grain shape (GLW) had negative effect on TGW at least for three QTL *QGlw.igdb-1A.2*, *QGlw.igdb-6A.1*, and *QGlw.igdb-7A.1*.

### Candidate genes underlying QTL through transcriptomic analysis

To detect dynamic profiles of genes involved in grain development, RNA sequencing (RNA-seq) was performed with whole spikes of two parents (KT1-1 and KT3-5) at four grain filling stages, 0 day after flowering (DAF), 7 DAF, 14 DAF, and 21 DAF. After filtering low-quality or adapter-contaminated reads, a total of 130.2 Gb clean data was harvested from 24 libraries of eight samples (two accessions × four developmental stages) and each with three biological replicates. After mapping against gene sets of the A genome of hexaploid wheat Chinese Spring (TGACv1.32, http://plants.ensembl.org/Triticum_aestivum), 40.38% reads covering 87.77% (28561 / 32539) of total genes have unique positions, which were subjected to further analysis.

The differential expressed (DE) genes between KT1-1 and KT3-5 were compared across four developmental stages using NOISeq (Tarazona *et al.*, 2011). A total of 4959 DE genes, including 2061 up-regulated and 2898 down-regulated genes, were identified with threshold of probability > 0.8 and |log_2_(fold change, FC)| > 1.0 (**Fig. S10a**). Through a Gene Ontology (GO) enrichment analysis, these genes were found involved in carbon fixation (GO:0015977, *P* < 10^-8^ at 7 DAF), amio acid metabolism (GO:0009069, GO:0009071, GO:1901606, *P* < 0.05), and regulations of nucleotide metabolism-related enzymes (**Fig. S10b**; **Table S11**). And they were mainly assigned to macromolecular complex (GO:0032991, *P* < 10^-4^), organelle part (GO:0044422, *P* < 10^-3^), specifically thylakoid related (GO:0009536, GO:0009534, GO:0031976land GO:0009579, *P* < 10^-4^) at 0 DAF, 7 DAF and 14 DAF, and cytoplasm (GO:0005737 and GO:0044444, *P* < 10^-2^) at 21 DAF in cellular components (**Fig. S10b**; **Table S11**). Regarding to molecular functions, structural molecule activity (GO:0005198, *P* < 10^-4^), enzyme activity that related with glucosyltransferase activity (GO:0046527, *P* < 10^-3^), hydrolase activity (GO:0016798, *P* < 10^-4^) and oxidoreductase activity (GO:0016491, *P* < 10^-4^) were dominant at 7 DAF, while enzyme, peptidase inhibitor and regulator activity related genes were prevailing at 21 DAF (**Fig. S10b**; **Table S11**). Furthermore, Kyoto Encyclopedia of Genes and Genomes (KEGG) enrichment analysis also revealed the divergence of the DE genes along with developmental stages (**Fig. S10c**; **Table S12**). The DE genes were highly enriched in the pathways related with energy metabolism (the photosynthesis, *Q* value < 10^-5^) and basic metabolisms at the early stage (0 and 7 DAF), while the genes at metabolisms of amino acid, fatty acid, hormone, nitrogen, and nutrients were mainly enriched at the middle stage (14 DAF). However, the protein processing and other possible biotic resistance-related metabolism (taurine, hypotaurine, and benzoxazinoid) expressed at the late stage (21 DAF).

To elucidate the candidate genes underlying QTL, DE genes on QTL regions were characterized through comparative transcriptional profiling along grain developmental process. Out of these mapped homologous genes, *SRS5* (*small and round seed 5*, TRIAE_CS42_5AL_TGACv1_377517_AA1247600.1) encodes an alpha-tubulin and regulated cell length and seed length in rice (Segami *et al.*, 2012). At the developmental stages critical for seed setting rate and the sink (grain) size, this gene had FPKM values of 547 and 650 at 0 and 7 DAF, respectively, in the high grain size parent KT3-5, while these were 227 and 257 in the low grain size parent KT1-1, respectively. This gene had overall up-regulation of 1.67 to 2.53-fold in KT3-5 at all the investigated stages (**Fig. 4**; **Table S13**). Regarding to the position on the genetic linkage map, *SRS5* was mapped onto the QTL region 5A-2 (142.8-164.3 cM), which contributed to GL, GW, GA and TGW (**Fig. 2**). The locus could explain 8.13% of the phenotypic variation for GL, and 7.23% for GW, further contributed to 22.29% (*R^2^*) of the phenotypic variation for GA (**Table 2**), implying that this locus affected grain size as in rice. Taken together, our data indicated that *SRS5* might be the candidate gene for this grain size QTL, affecting ~6.22% GA and ~8.78% TGW (**Table S6**).

**Fig. 4.**
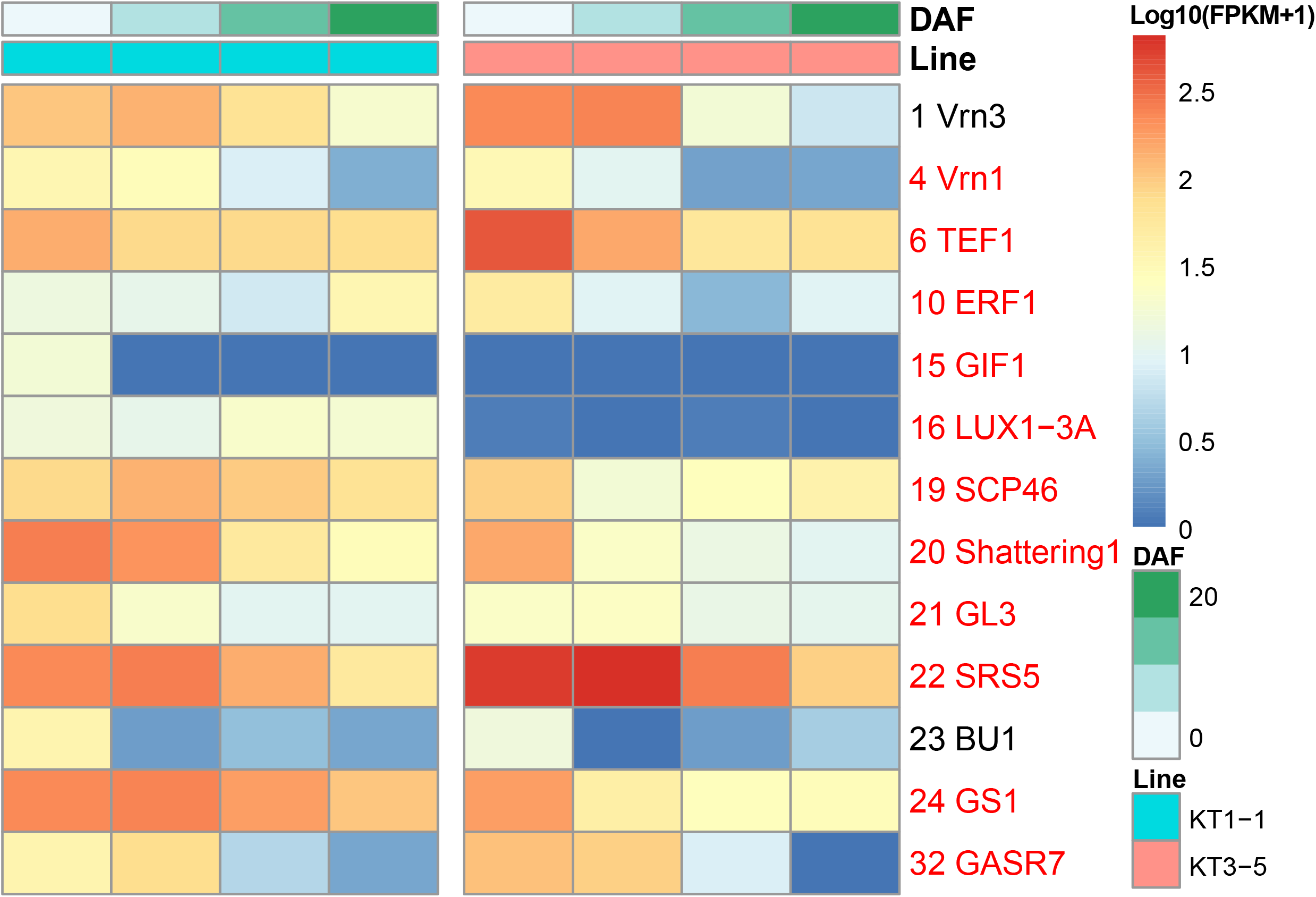
Transcriptional profiles of genes mapped to QTL regions in two parental lines. Only genes differentiated expressed in at least one developmental stage (probability > 0.7) were retained, while genes with names in red were probability > 0.8 from NOISeq. The log_10_(FPKM+1) transformed data were plotted.

*TEF1* is one of transcript elongation factors and affected tilling number in rice but increased grain size in wheat (Zheng *et al.*, 2014). This gene was mapped onto two genomic regions 1A-3 (200.8-224.3 cM) and 7A-1 (62.3-70.4 cM). The copy on 1A-3 showed higher expression levels at 0 DAF and 7 DAF in KT3-5 than in KT1-1, while it had similar levels between them at late developmental stages (14 to 21 DAF) (**Fig. 4**). The cultivated allele of QTL on 1A-3 could increase about 5.04% TGW (**Table S6**). Moreover, *TEF1* showed relative high expressions (FPKM > 57) in grain (**Table S13**), which might confer to phenotypic variations of grain (sink) size of developing seed. However, another copy on 7A-1 expressed increasingly along with grain development stages and reached to maximum at 21 DAF, which was similar with its expression profiling in common wheat (Zheng *et al.*, 2014). Nevertheless, no significant differences of expression patterns between two parents were observed across grain filling stage, consistent with no QTL for TGW on 7A-1 (**Table 2**).

The gene encoding ADP-glucose pyrophosphorylase (*AGPase*) large subunit was mapped to the genomic region 1A-3 (200.8-224.3 cM) based on homology analysis, which was a rate-limiting enzyme to catalyze the formation of ADP-glucose (ADPG), the substrate for starch biosynthesis (Georgelis *et al.*, 2007). This gene (*AGPLcyto*, ID17) was highly expressed with FPKM values of 398 and 246 at the middle and late stages of grain development, respectively (**Fig. 5**; **Table S13**), which is crucial for grain filling (Yang *et al.*, 2004). Furthermore, the gene of AGPase small subunit (*AGPScyto*, ID18) had similar pattern with high expression levels (FPKM reaching to 1194 at 14 DAF). These two genes were differentially expressed between the two parental lines at 7 DAF for *AGPLcyto* and 14 DAF for *AGPScyto* (probability > 0.7), respectively (**Fig. 5**; **Table S13**). To confirm the gene location and elucidate the causality of expression variation, *AGPL* in einkorn wheat was sequenced and several variations were detected along both the promoter and genic regions between two parental sequences, including InDel and SNP (**Fig. S9**). A polymorphic marker based on one InDel at intron I was co-localized with genomic region 1A-3 on this genetic map, further confirming the reliability of its homology-based mapping. Our data indicated that variations of expression levels for its two subunits determined the formation of ADPG, and further affected the starch accumulation and even grain weight.

**Fig. 5.**
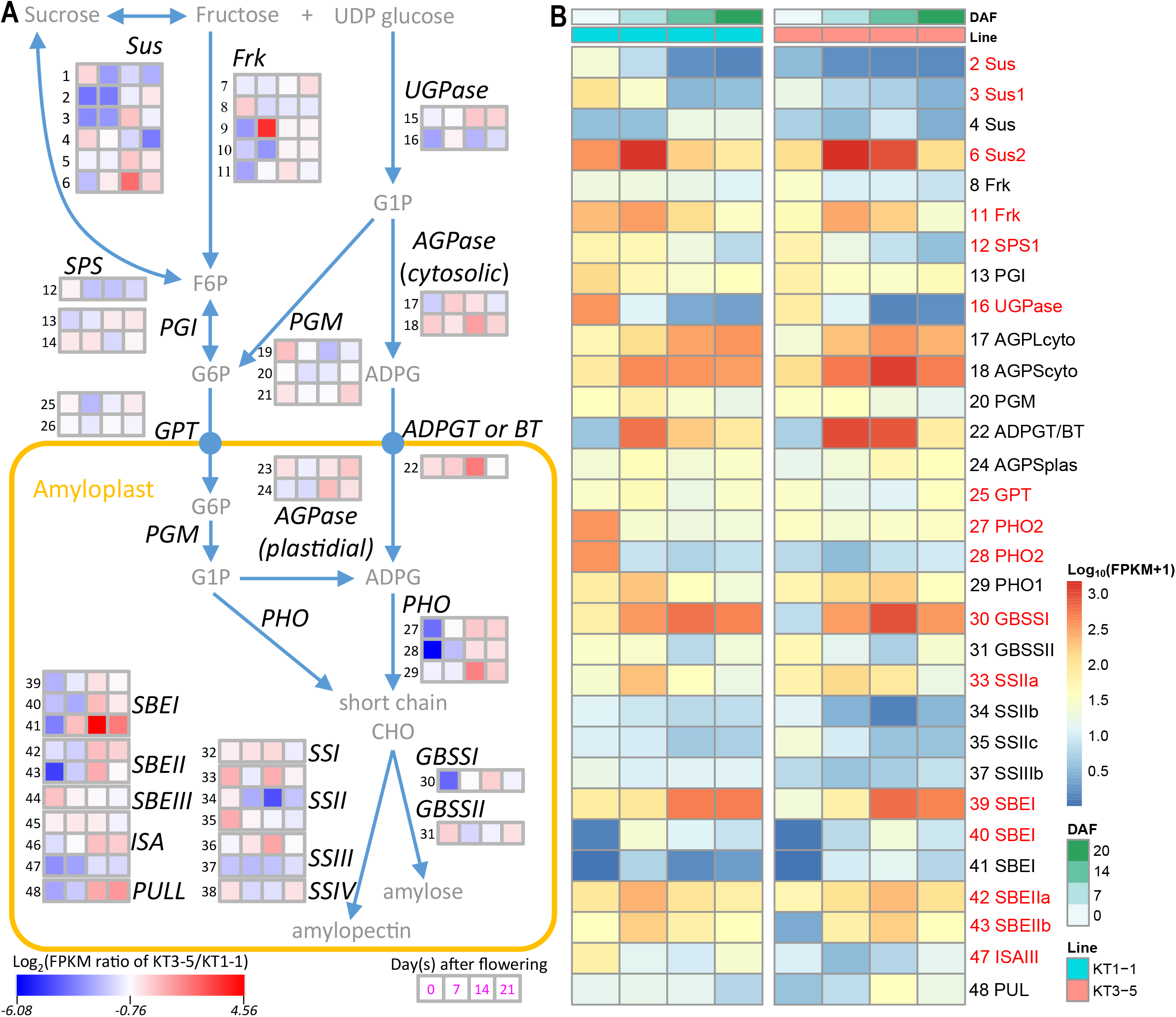
Fold changes and expression patterns of starch biosynthesis genes across four grain developmental stages between two parental lines. Only genes differentiated expressed in at least one developmental stage (probability > 0.7) were retained, while genes with names in red were probability > 0.8 from NOISeq. (a) Wheat starch biosynthesis pathway. The Log_2_ fold change of KT1-1 *vs* KT3-5 in FPKM at four developmental stages were ploted in heatmp. (b) Heatmap of the expression profiles of the starch biosynthesis pathway genes in grains of two parental lines, KT1-1 (left) and KT3-5 (right). The log_10_(FPKM+1) transformed data were plotted in (b).

To further investigate the starch biosynthesis pathway involved in the grain development, forty-eight genes in the pathway were retrieved from CS cDNA database based on previous information (Krasileva *et al.*, 2017), and the expression patterns of their homologs in both parental lines were compared. In total, thirty-one genes (65% of total genes) were significantly differentially expressed at least one stage between KT1-1 and KT3-5 with probability > 0.7 (**Fig. 5**). Out of these genes, the restricted enzyme gene, *ADPGT* (*ADPG Transporter*) or *BT1* (*Brittle1*), transporting the substrate for starch biosynthesis ADPG from cytoplasm to amyloplast in cell (Sullivan *et al.*, 1991), was highly expressed in KT3-5 (FPKM > 800) with fold changes of 1.67 and 4.86 (probability > 0.7) for 7 DAF and 14 DAF, respectively. Another gene, *Starch Phosphorylases 1* (*PHO1*, ID 29), which is responsible for the conversion of ADPG to the precursor for starch biosynthesis by starch synthase (SS, EC 2.4.1.1), was expressed in KT3-5 above 4-fold changes than in KT1-1 at 14 DAF. Moreover, *Sucrose synthase* (*Sus*, ID 6) had much higher expression level, especially at late developmental stages, which could compensate the expressions of another five low-abundance *Sus* copies (ID 1~5) (**Fig. 5**; **Table S13**). The *UDP-glucose pyrophosphorylase* (*UGPase*, ID 15), *Starch branching enzyme IIa* (*SBEIIa*, ID 42) and *SBEIIb* (ID 43) showed similar patterns as well. Though several DE genes were expressed highly in KT1-1 at 0 DAF or 7 DAF, for example, *Sus* (ID 2 and 3), *Frk* (ID 11), *UGPase* (ID 16), *PHO2* (ID 27), *PHO2* (ID 28), *GBSSI* (ID 30), *SBEI* (ID 39), *ISAIII* (ID 47) and *PUL* (ID 48), most of these genes were less down-regulated or even up-regulated in KT3-5 at late developmental stages, which were a very critical period for starch accumulation (Yang *et al.*, 2004). In developing seeds, these rate-liming functional enzymes are very important for starch synthesis, and differentially expressed of relevant genes would affect the starch accumulation and further grain development.

In summary, through QTL analysis, homologous gene mapping and transcriptomic analysis, 44 of total 80 genes on QTL regions or in starch biosynthesis pathway were mapped on nine QTL regions and showed differentially expressed in two parental lines (**Table S13**). Our data demonstrated that the expression patterns of several functional genes were consistent with allelic effects of the related QTL, which implied that they were the candidate genes underlying QTL. The candidate functional genes associated with grain (sink) size and starch biosynthesis were considered as important components for grain size and starch accumulation in developing grains, in turn for grain weight, explained most of the phenotypic variations ranging from 52.3% for GLW to 66.7% for TGW (**Table S9**). Furthermore, the phenotypic values of the related traits significantly increased as the numbers of the positive QTL alleles accumulated (**Fig. S8**), which could be exploited to fine-tuned grain size and weight. Thus, our data revealed the complex genetic architecture of gain size on a genome-wide scale, elucidated QTL and their underlying genes through genetic mapping and transcriptional profiling, which were an essential for identifying the candidate genes for grain size and weight and could assist marker-based selection in wheat breeding improvement.

## Discussion

Grain size, as a complexity trait, is still less understood in wheat. In the respect of diploid nature of genome and richness of natural diversity of grain size, we revealed genetic architecture of grain size using an einkorn wheat RIL population. In this study, the RAD-seq, combining NGS and restriction enzyme digestion to reduce the genome complexity, was explored to provide the genetic polymorphisms at a genome scale for the development of a high-density genetic map integrated with previously einkorn wheat map (Yu *et al.*, 2017). This high-density map contains 10876 evenly distributed genetic markers, had 1551 unique positions, and covered > 97% physical maps of wheat and barley genomes, demonstrating its good quality for einkorn wheat genetic and genomic researches. In particular, the comparative genomics through mapping SNP markers from genetic map could not only aid to examine the syntenic blocks with genomes of wheat relatives, but also provide the genomic sequence information for dissecting interesting regions and for revealing structural variations of chromosomes between different genomes.

### The grain size variations in einkorn wheat were elucidated by phenotypic and genome-wide QTL analysis

In this study, the RAD-seq derived high-density genetic map was explored to access the genetic architecture of grain size. Phenotypic analysis demonstrated that grain size traits were under complex genetic control, the strong correlations were identified between these traits. Two PCs captured grain size and shape, respectively, explaining > 97% of the phenotypic variations. The genome-wide QTL analysis were further conducted and identified a total of 17 genomic regions that contributed to grain size and weight. Five genomic regions on chromosome Tm5A were associated with most surveyed traits, which coincided with three meta-QTL on chromosome Ta5A of common wheat in a meta-analysis with six double haploid populations (Gegas *et al.*, 2010). The mapping interval 1A-2 containing QTL for GA and TGW was co-localized with the meta-QTL MQTL2, which was linked with *Glu-1A* on Ta1A (Gegas *et al.*, 2010). Moreover, two QTL regions distributed on Tm7A were associated with all the screened traits, but in common wheat, only QTL for GL, GLW, GA, TGW were detected from two individual populations and no meta-QTL was reported (Gegas *et al.*, 2010). In addition, the distribution breadth of QTL (the number of chromosomes and genomic regions on which QTL were detected) identified from this individual RIL population of einkorn wheat is wide, for which two importantly possible reasons were relatively broad polymorphisms between two surveyed parental lines and high genomic coverage of the genetic map. Thus, it demonstrated that the present RIL population for einkorn wheat harbored more genetic diversities than common wheat, although the principal components extracted from phenotypic dataset were similar between them, but this RIL population have smaller background-specific effects in determining the genetic architecture not as in hexaploid wheat population (Gegas *et al.*, 2010).

### Novel QTL were detected in einkorn wheat through comparing with tetraploid and hexaploid wheat populations

QTL for grain size have been widely studied in wheat, and they were detected all chromosomes of tetraploid and hexaploid wheat (Brinton *et al.*, 2017; Cheng *et al.*, 2017; Gegas *et al.*, 2010; Golan *et al.*, 2015; Kumar *et al.*, 2016; Maphosa *et al.*, 2014; Peleg *et al.*, 2011; Prashant *et al.*, 2012; Rasheed *et al.*, 2014; Russo *et al.*, 2014; Tsilo *et al.*, 2010; Williams and Sorrells, 2014; Wu *et al.*, 2015). However, very limited information is available for QTL analysis of grain size and weight in einkorn wheat. Based on syntenic regions between einkorn wheat and hexaploid wheat (**Fig. 3**) and marker information on Chinese Spring RefSeq v1.0 genome (https://urgi.versailles.inra.fr/jbrowseiwgsc/gmod_jbrowse/), QTL regions overlapped markers from other studies were considered as common QTL regions, otherwise novel QTL regions. In our study, five of seventeen QTL regions were newly detected, including 1A-3, 5A-4, 5A-5, 6A-2, and 7A-3. Three QTL regions, 3A-2, 5A-2 and 7A-1, involving 12 QTL for grain size and weight, should be affected by heading date under the control of three genes *TmLUX1*, *Vrn1* and *Vrn3*, compared our previous findings (Yu *et al.*, 2017). The other 12 QTL regions overlapped with QTL or markers from studies in tetraploid and hexaploidy wheat (Brinton *et al.*, 2017; Cheng *et al.*, 2017; Gegas *et al.*, 2010; Golan *et al.*, 2015; Kumar *et al.*, 2016; Peleg *et al.*, 2011; Russo *et al.*, 2014; Wang *et al.*, 2009; Wu *et al.*, 2015). For example, a TGW QTL detected in tetraploid wheat population linked with *wPt-7053* (Peleg *et al.*, 2011), which locates on 676.40 Mb of 7A, and similarly, QTL-27 (Kumar *et al.*, 2016) and *QGl.cau-7A.3* (Wu *et al.*, 2015) from hexaploid wheat were located 674.27-705.13 Mb and 671.42-679.96 Mb, respectively, corresponding to 7A-2 region (670.94-693.33 Mb) from this study. However, 7A-2 was detected to be associated with GL and GLW in this study, while in hexaploid wheat this locus affected GL/GW/GA/TGW and GL only, respectively (Kumar *et al.*, 2016; Wu *et al.*, 2015). Our data indicated that einkorn wheat had similar genetic basis but divergent functions of some locus of regulating grain size with tetraploid and hexaploid wheat.

### The candidate genes underlying QTL were predicted based on comparative genomics and transcriptomics

The underlying candidate genes of genetic loci could help to understand the morphogenesis, and to provide diverse alleles for breeding improvement in common wheat. Last two decades has witnessed the characterization of a number of causative genes for grain size and weight in crops (Li and Yang, 2017). In this work, by comparative analysis and transcriptomic profiling with RNA-seq, several genes have been proved to be the candidate genes for grain size in einkorn wheat. Those mapped homologous genes were differentially expressed at early to middle stages for expanding grain volume before initiating grain filling in developing seeds. One copy of a transcript elongation factor (*TEF*) gene on Tm1A was highly expressed in the KT3-5 with big seeds at 0 DAF and 7 DAF. Overexpression of *TEF* of hexaploid wheat could enhance the grain length in Arabidopsis, and *TEF1* was significantly associated with grain length and width, and TGW in wheat through haplotype analysis (Zheng *et al.*, 2014). Moreover, the homolog of *SRS5* of rice showed successive higher expression in cultivated einkorn wheat KT3-5 across all developmental stages. This gene encoding alpha-tubulin protein mainly expressed on young panicle and regulated cell elongation and seed length (Segami *et al.*, 2012). Mutant *SRS5* (Os11g0247300) with an amino-acid substitution reduced seed length 1.38 mm by decreasing cell and lemma length. The wild type *SRS5* could partially rescue mutant phenotype in transgenic plants (Segami *et al.*, 2012). Thus, these genes mapped on QTL regions were differentially expressed at the start of grain development and should be considered as main determinant factors for grain size. Furthermore, starch accumulation was deliberated critically to grain filling in wheat (Yang *et al.*, 2004), whose pathway has been well-exemplified (map00500 in KEGG, http://www.kegg.jp). Five enzymes, Sus, AGPase, AGPGT (BT1), SS and SBE were characterized to be critical for this process, and their encoding genes play the important roles in formation of UGPG (the first step in the conversion of sucrose to starch) and ADPG (the substrate of starch), transferring ADPG into amyloplast from cytoplasm and yielding the end starch(Sullivan *et al.*, 1991; Yang *et al.*, 2004). In this work, differential expression of these rate-limiting enzyme encoding genes were observed at middle to late stages of grain development. The *AGPL* co-localized with 1A-2, involved QTL for TGW, GW and GLW (**Fig. 2**), and the negative allele from KT1-1 can decrease 1.20 g TGW accounting for 7.8% to 15.4% of parental phenotypic variations across the surveyed environments (**Table 2**; **Table S1**). Further several genomic variations including SNPs and InDels were observed in this gene involving promoter and genic regions, implying that transcriptional levels differentiated between two parents might due to these variations (**Fig. S9**). Alleles for *Sus* and *AGPL* have closely association with grain weight in common wheat, mainly contributed by variations on transcript levels, which affected about 3-5 g and 2-4 g TGW, respectively (Hou *et al.*, 2017; Jiang *et al.*, 2011). Overall, transcriptional profiling analysis of 48 genes in starch biosynthesis pathway were investigated to elucidate expression patterns in developing seeds of einkorn wheat in this work. Thirty-one genes were expressed differentially at different grain development stages, especially the aforementioned rate-limiting enzyme genes (**Fig. 5**). Therefore, our data indicated that different expression patterns of these pathway genes together might contribute the final grain weight by affecting starch accumulation across grain filling. Furthermore, we developed molecular markers for these related genes, for example *AGPL*, *NAL1-2A* (ID 8), *GS1a* (ID 24), *GASR7* (ID 32), and flowering pathways genes (*Vrn1*, *Vrn2* and *Vrn3*), which will facilitate in marker-assisted wheat breeding endeavors (**Table S7**).

### Genes in flowering pathway affect the grain size and weight

Several genes in flowering pathway, such as *Vrn1*, *Vrn2*, *Vrn3*, and *TmLUX1* were mapped onto our genetic linkage map (**Fig. 2**). These genes have been positional cloned and functioned in wheat flowering pathway (Gawronski *et al.*, 2014; Yan *et al.*, 2006; Yan *et al.*, 2003; Yan *et al.*, 2004), and QTL analysis identified them as candidate genes underlying QTL for heading date (Yu *et al.*, 2017). In this study, they were mapped or linked to QTL regions involving grain size traits, *Vrn1* on 5A-2, *Vrn2* closely linked to 5A-4, *Vrn3* on 7A-1, and *TmLUX1* on 3A-2 (**Fig. 2**). As expected, these reproductive development-related genes have further affected the final performance in grain size and weight. The cultivated einkorn wheat alleles of QTL at *Vrn1* and *Vrn2* loci have positive effects on all related traits (mainly grain size related, GL, GW, GA, and TGW), while the wild type allele on QTL at *Vrn3* locus showed positive effects on grain shape related traits (GL and GC) and promoted flowering (Yu *et al.*, 2017). This may imply that the vernalization requirement is an important domestication syndrome in einkorn wheat, while heading or flowering time were not selected for their influence on the adaptation to different growth environments in einkorn wheat (Bullrich *et al.*, 2002; Lewis *et al.*, 2008; Snape *et al.*, 2001).

### The untapped alleles for grain weight were identified in wild einkorn wheat

In this work, we identified seven QTL within three QTL regions (7A-1, 7A-2, and 7A-3) on chromosome Tm7A, of which the wild einkorn wheat has positive alleles for all the grain size related traits. They contributed to grain length and width in a relative large proportion, for example, *QGl.igdb-7A.2* on 7A-2 increasing 0.19 mm grain length and explaining 15.5% of the phenotypic variation of GL, *QGw.igdb-7A.1* co-locating with both *QGa.igdb-7A.1* and *QTgw.igdb-7A.1* on 7A-3 to increase the 0.55 mm^2^ grain area and 1.29 g thousand-grain weight (**Table S6**; **Table 2**). It indicates that the remaining superior alleles of genes controlling improvement of grain size might be ignored by preliminary selection for domestication at ~10000 year ago, and some alleles that contribute to a wide adaptation of wheat to different environments (such as *Vrn3*) were left in wild einkorn wheat. In addition, a gene *Shattering* (*Sh1*) encoding a YABBY transcription factor, have been reported in rice, sorghum and maize for shattering phenotype, one of important domestication syndrome (Lin *et al.*, 2012). This gene was highly expressed in wild einkorn wheat with brittle rachis on field natural state, which implied that this gene might be one another genetic locus determining brittle rachis and/or was selected in domestication history of einkorn wheat, as *Q* and *Btr* loci in wheat and barley (Avni *et al.*, 2017; Dubcovsky and Dvorak, 2007; Pourkheirandish *et al.*, 2015). Thus, we speculate that there is a part of dominant genes or alleles to enlarge the grain size and further to increase the grain weight, could be characterized from natural wild einkorn wheat for the candidate potentials.

## Acknowledgements

We thank Chikako Shindo and Tetsuo Sasakuma (Kihara Institute for Biological Research/Graduate School of Integrated Science, Yokohama City University, Japan) for providing experimental materials used in this study. This work was supported by National Key Research and Development Program of China (2016YFD0101802) and the National Natural Science Foundation of China (31571643).

## Author’s contributions

AZ, CL conceived and supervised the study; KY, DL, YC, DW, WLY, WY, LY, CZ, SZ and JS conducted the research and analyzed the data; KY and DL planed and conducted the construction of RAD-seq libraries; KY, DW, WLY, and JS collected phenotypic data; KY analyzed the genotypic and phenotypic data; KY collected samples for transcriptomic analysis; KY, YW, LY, CZ and SZ contributed to analyses of transcriptomic data; KY, DL, and AZ prepared the manuscript. All authors discussed the results and commented on the manuscript.

## Supplementary data

**Fig. S1** Frequency distribution of phenotypic data in five environments for six quantitative traits of parents and RILs.

**Fig. S2** Principal component analysis revealing a morphometric model for variation in grain morphology in einkorn wheat RIL population.

**Fig. S3** Chromosomal 4AL/5AL translocation and 4A pericentric inversion in hexaploid wheat revealed by comparing barley and hexaploid wheat genomes through the mapped SNP markers from einkorn wheat. (a) The einkorn wheat genetic map was compared with barley genome. (b) Physical positions of SNP markers on barley genome (X-axis) was compared with positions on Chinese Spring A genome physical map (Y-axis), which elucidated 4AL/5AL translocation event in A genome of hexaploid wheat and einkorn wheat, and pericentric inversion on 4A in hexaploid wheat. (c) Genomic coverages of the physical maps of hexaploid wheat and barley spanned by the einkorn wheat genetic map.

**Fig. S4** Genome-wide LOD profiles for six investigated traits across five environments. (a) LOD profile; (b) phenotypic variation explained by QTL; (c) additive effects by KT1-1 allele.

**Fig. S5** Physical lengths of homologous regions corresponding to sixteen QTL regions.

**Fig. S6** Polymorphism of *Vrn3* in einkorn wheat. A 6-bp InDel was detected at −195 upstream of the promoter region of *Vrn3*, and molecular marker (Vrn3-InDel-F1: TGAACTGGTCTGGACATGGA in red and Vrn3-InDel-R1: GGAGCAAGCAGCAGAGGTTA in green) was developed around this variation, producing 162 bp at KT1-1 and 168 bp at KT3-5.

**Fig. S7** Genetic overlaps between grain size related traits in the einkorn wheat RIL population. Distribution of 42 unique QTL at seventeen QTL regions for six investigated traits across five environments.

**Fig. S8** Phenotypic variations affected by positive number of allele for six quantitative traits using the average phenotypic data. (a) Mean phenotypic values for each trait. (b) Phenotypic values of six trait from each environment.

**Fig. S9** Polymorphism of *AGPL* in einkorn wheat. An InDel were detected at Intron I of *AGPL*, and molecular marker (AGPL-F1: CTCCAGGAGGATGTGCAAC in green and AGPL-R1: CAGAGATGCTAACATAACAGAGTG in red) was developed around this variation, producing 156 bp at KT1-1 and 165 bp at KT3-5. Positions of exon and intron were denoted with brown boxes.

**Fig. S10** GO and KEGG enrichment analysis of differential expressed (DE) genes identified from four developmental stages of KT1-1 *vs* KT3-5. Numbers in red and black denoted up and down-regulated genes, respectively. (a) Overlaps of DE genes from four developmental stages. (b) GO enrichment analysis of DE genes, * *P* < 0.05, ** *P* < 0.01. (c) KEGG enrichment analysis of DE genes, * *P* < 0.05 (left) or *Q* < 0.05 (right), and pathway names in red and green denote the *Q* < 0.05 and > 0.05, respectively.

**Table S1** Summary statistics and broad-sense heritability of six grain size related traits.

**Table S2** Correlation coefficients between GL, GW, GLW, GA, GC, and TGW in the RIL population in six environments.

**Table S3** Principal components analysis and correlation with phenotypic data. Probability loadings of principal components identified in four environments (a) and Pearson’s correlation coefficients of two principal components with phenotypic data in each environment (b).

**Table S4** Barcode and sequencing information of RAD-seq.

**Table S5** The high-density genetic linkage map of einkorn wheat.

**Table S6** Individual QTL detected with CIM method from five environments using the high-density SNP map of einkorn wheat.

**Table S7** Candidate genes mapped to QTL region based on homologous analysis with hexaploid wheat, barley and rice.

**Table S8** Polymorphic markers of functional genes.

**Table S9** Phenotypic variation explained by the detected QTL estimated using analysis of variance (ANOVA) results from a simple model and multiple regression analysis. Each of the detected QTL are represented by their peak markers.

**Table S10** Multiple comparison test of phenotypic variations affected by positive number of allele for six quantitative traits across all five environments. One-way ANOVA followed by Tukey’s Honestly Significant Difference (HSD) test was performed, and different letters were assigned to significantly different groups (P < 0.05). NA for not available.

**Table S11** GO enrichment test of DE genes identified in einkorn wheat grains from four developmental stages.

**Table S12** KEGG enrichment test of DE genes identified in einkorn wheat grains from four developmental stages.

**Table S13** Expression profiles of genes involving in starch biosynthesis pathway and candidate genes mapped to QTL region based on homologous analysis with hexaploid wheat, barley and rice.

## Abbreviations

ANOVA: analysis of variance
CIM: composite interval mapping
cM: centiMorgan
GA: grain area
Gb: giga base pairs
GC: grain circumference
GL: grain length
GLW: grain length / width
GW: grain width
InDel: insertion deletion
LOD: Logarithm of the odds
Mb: mega base pairs
QTL: quantitative trait locus or loci
RAD-seq: restriction site-associated DNA sequencing
RIL: recombinant inbred line
RNA-seq: RNA sequencing
SNP: single-nucleotide polymorphism
TGW: thousand-grain weight

